# Policy regularization as a unifying theory of the striatal division of labor in learning

**DOI:** 10.64898/2026.09.10.750654

**Authors:** Cheshta Bhatia, Samuel J. Gershman, Lucy Lai

## Abstract

Learning novel behaviors requires balancing previously learned actions with the ability to flexibly adapt to changing reward contingencies. This trade-off is well documented in the division of labor between dorsolateral striatum (DLS), which promotes selection of cached, history-dependent actions, and dorsomedial striatum (DMS), which supports flexible learning as reward contingencies change. Here, we propose policy regularization as a general computational principle for understanding this division. Capacity-limited agents face a fundamental trade-off between maximizing reward and minimizing the cost of deviating from a default policy that caches frequently used action transitions. We formalize this trade-off as a KL-regularized reward objective in which a flexible controller (DMS) incurs a cost for diverging from a history-dependent default (DLS). The resulting optimal policy is a combination of a reward-driven action value, continuously updated by DMS, and a default policy conditioned on action history, cached by DLS. In practice, DLS consolidates the action transitions shaped by DMS’s reward-driven value learning, so DLS preserves behaviors that were once optimal even when reward contingencies change—producing robust but inflexible action selection and effectively “regularizing” DMS-driven learning. We show that a single model—with a single set of learning and lesion parameters, shared across paradigms—provides a unifying explanation for the functional organization of the striatum, reproducing canonical DLS–DMS dissociations in outcome devaluation, serial spatial reversal, skilled action sequencing, and motor sequence execution tasks.

**Author Summary:** How does the brain balance old, well-practiced behaviors with the flexibility to adapt when the world and reward contingencies change? Two neighboring regions of the dorsal striatum appear to divide this work: lesions of the dorsomedial striatum impair learning of new or changed action–outcome contingencies, whereas lesions of the dorsolateral striatum impair execution of habits and learned action sequences. We propose a theoretical framework in which a single principle—maximizing reward while minimizing the cost of deviating from previously cached action patterns—explains this division of labor: the dorsomedial striatum learns reward-driven action values that support flexible, goal-directed behavior, while the dorsolateral striatum caches recurring action transitions regardless of current reward. A single model reproduces canonical lesion dissociations across four behavioral paradigms (reward devaluation, reversal learning, skilled lever-press sequences, and cued versus automatic movement execution), offering a normative explanation for why these two subregions exist and compete, and providing a potential circuit implementation underlying learning.

## 1 Introduction

Intelligent behavior requires both automatic, well-rehearsed action patterns and the capacity to adapt when circumstances change. Cached, history-dependent behaviors support rapid action selection in familiar contexts, while flexible control adapts to novel demands—but the two pull in opposite directions: automaticity sacrifices flexibility, and flexibility consumes cognitive resources.

In reinforcement learning (RL), capacity-limited agents balance these demands by maximizing reward while minimizing the information cost of complex policies [Still and Precup, 2012, Tishby and Polani, 2011, Parush et al., 2011, Rubin et al., 2012]. This realizes a form of *policy compression*: simplifying policies to reduce cognitive cost [Gershman, 2020, Lai and Gershman, 2021, 2024]. Compressed policies are also reusable: once a policy is consolidated, the same actions can be deployed across contexts without recomputation, supporting fast and automatic action selection. The trade-off is flexibility: when the environment changes, a compressed policy may no longer recover the optimal action. How do capacity-limited agents trade reusable compressed policies against flexible action selection?

Policy regularization is a method that uses a “default” policy to regularize a “control” policy, reducing the information cost of the agent’s behavior [Geist et al., 2019]. In particular, Kullback–Leibler (KL) regularization penalizes the control policy for being too “far” from the default, as measured by the KL divergence [Tirumala et al., 2019, Galashov et al., 2019, Teh et al., 2017]. The default policy is learned alongside the control policy and provides a cost-effective mechanism for caching and reusing frequent behaviors. To provide an intuitive example, consider a chef who specializes in spaghetti and meatballs: after making it many times, the action sequence becomes highly practiced and can be reused with little effort. Preparing a similar noodle dish, such as ramen, may therefore be relatively low-cost because it shares some familiar actions (e.g., boiling noodles), whereas making sushi requires a more distinct sequence of actions and thus incurs greater cognitive cost.

KL regularization has been successful in solving challenging RL problems that involve complex and diverse tasks by improving exploration and generalization [Schulman et al., 2015, Levine, 2018, Abdolmaleki et al., 2018], and can be viewed as an implementation of policy compression that penalizes deviation from a default policy. Here, we use KL regularization as a normative framework to understand how biological agents arbitrate between reusing cached behavior and adapting to change.

Biological organisms face the same tension between automaticity and flexibility [Del Giudice and Crespi, 2018]. For example, a mouse executes stereotyped predatory routines [Yu et al., 2021], but must update them by incorporating external environmental factors in the search [Gire et al., 2016, Kőszeghy et al., 2025]. Within the brain, one well-documented example of how neural circuits implement this balance is found in the functional specialization of basal ganglia circuits. We posit that the DLS–DMS division of labor arises from this fundamental trade-off. Specifically, we hypothesize that DLS encodes the default policy *π*_0_, supporting automatic, history-dependent action selection, while DMS learns the reward-driven action values *Q* that adapt to current reward contingencies. Their interaction, governed by the relative weighting of *π*_0_ and *Q* in the control policy, determines whether behavior is habitual or flexible (Figure 1). Under this framework, DLS effectively regularizes DMS by biasing behavior toward previously consolidated action patterns, reducing the complexity and information cost of control. We demonstrate that a single KL-regularized model, with consistent lesion implementations across paradigms — ablating the default policy for DLS and slowing value learning for DMS — reproduces canonical DLS–DMS dissociations in four behavioral paradigms: outcome devaluation [Yin et al., 2004], serial spatial reversal [Castañé et al., 2010], skilled action sequencing [Turner et al., 2022], and motor sequence execution [Mizes et al., 2023]. These results provide a normative rationale for the functional organization of the striatum, suggesting that the DLS–DMS division of labor reflects a principled trade-off between minimizing behavioral cost and maximizing reward.

**Figure 1:**
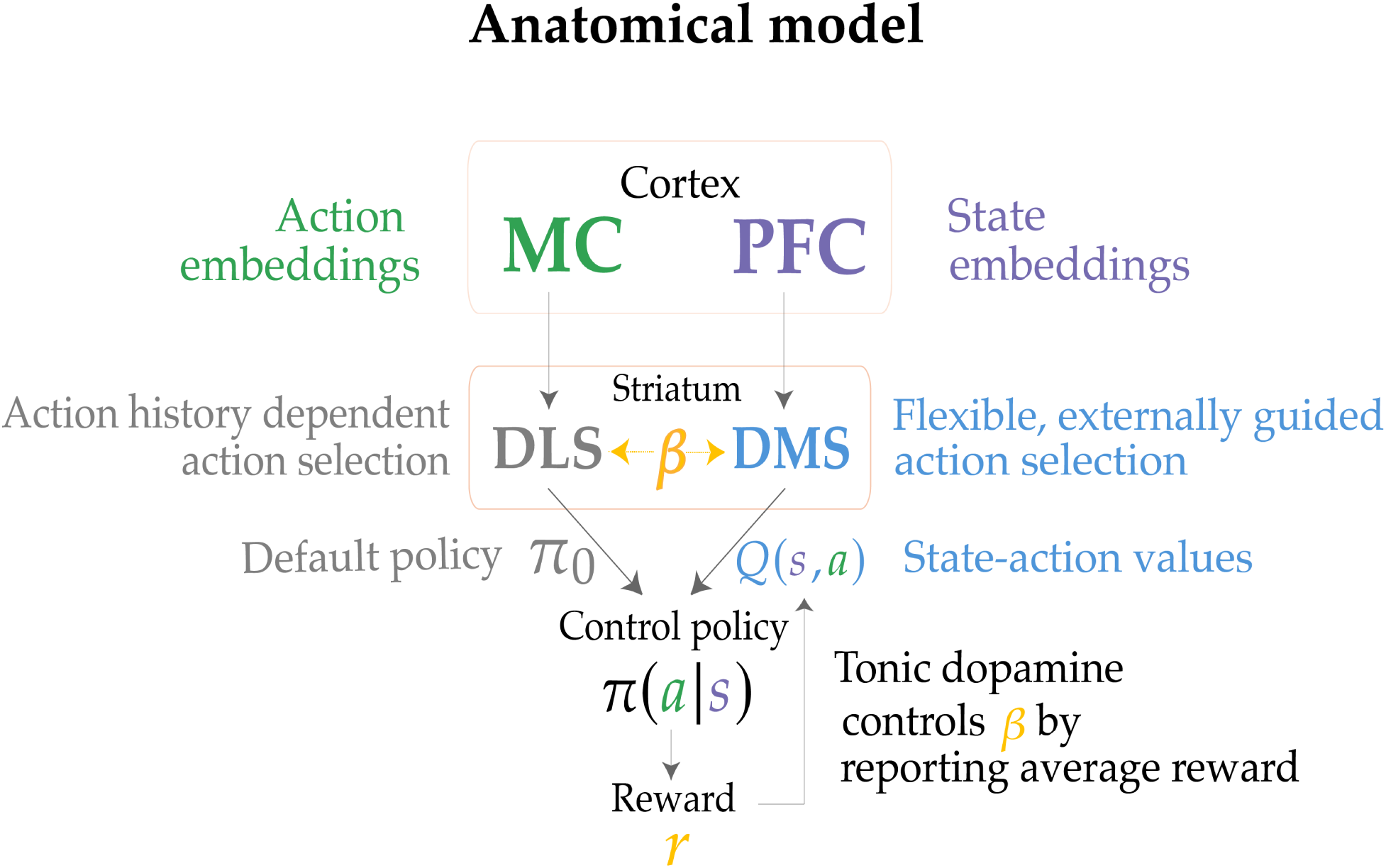
Policy regularization model of the DLS–DMS division of labor. Anatomical mapping: action-related (MC) and state/context-related (PFC) cortical representations feed DLS and DMS, respectively, whose outputs converge to form the control policy.

## 2 Materials and Methods

### 2.1 Theoretical framework: Policy Regularization

#### 2.1.1 Model setup and KL regularized objective

We model behavior as a Markov Decision Process (MDP) M = (S, A, *P*, *r*, *γ*), where S is the state space, A is the action space, *P*(*s*^′^ | *s*, *a*) is the transition function (the probability of transitioning to state *s*^′^ ∈ S after taking action *a* ∈ A in state *s* ∈ S), *r* (*s*, *a*) is the reward function, and *γ* ∈ [0, 1] is the discount factor. The state *s* represents the agent’s current situation that action selection depends on — it includes external sensory information (such as visual, auditory, or olfactory contextual cues that bias decision making) and internal states (such as agent’s motivational state). Throughout, we adopt a consistent convention: *states* correspond to the external/internal cues that specify which action is currently appropriate and *actions* correspond to the animal’s responses (e.g., pressing a particular lever or poking a particular hole). The number of states in each paradigm therefore reflects its cue structure, not the number of response options. In cue-guided tasks the state indexes the current cue—five step-cues in the action sequencing task [Turner et al., 2022] and three cues in the motor sequence execution task [Mizes et al., 2023] —whereas in uncued tasks the external state is latent. When no discriminative cue is ever present, as in the spatial-reversal task [Castañé et al., 2010], the external state collapses to the presence of levers and the correct choice must be inferred from reward feedback rather than read from a cue.

We treat the agent as model-free: it does not learn or use *P* explicitly, and instead learns values directly from experienced rewards via temporal-difference updates. We note that the framework could easily be extended to model-based settings by equipping the agent with an estimate of a transition function for planning, while retaining KL regularization as a cost for deviations from the default policy. The dopaminergic signal that drives value learning in our model may also support such model based learning [Akam and Walton, 2021]. The agent maintains two policies:

- A **control policy** *π*(*a_t_* | *s_t_*) used to select actions.
- A **default policy** *π*_0_(*a_t_* | *a_t_*_−1_), a distribution over actions conditioned on the previous action and independent of the external state. The default policy caches frequently used action transitions, providing a low-cost solution for action selection.

Following the default policy is the agent’s low-cost baseline; deviating from it requires effort. Selecting actions that diverge from *π*_0_ requires the agent to override cached transitions and incorporate external environmental information, which is information-theoretically expensive [Lai and Gershman, 2021]. We adopt the KL divergence as a measure of how much the control policy *π*(| *s_t_*) deviates from the default policy *π*_0_(· | *a_t_*_−1_):

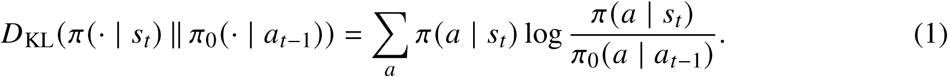

*D*_KL_ is non-negative, equals zero when *π* = *π*_0_, and grows as the two distributions diverge. Compressing the policy means keeping this divergence small (i.e., selecting actions that do not deviate too far from the default *π*_0_). The agent’s goal combines two competing requirements:

- Maximize expected discounted reward, E*_π_* [Σ*_t_* *γ^t^r* (*s_t_*, *a_t_*)].
- Minimize the cumulative KL divergence of *π* from *π*_0_.

Capacity-limited agents are therefore motivated to learn compressed policies that stay close to the default *π*_0_. We formalize this trade-off via the following KL-regularized reward objective:

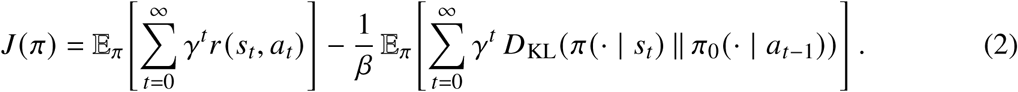

The inverse temperature *β* governs how much state-specific information the agent is willing to encode at any moment. Large *β* makes deviation cheap, recovering near-greedy reward maximization, while small *β* makes deviation expensive, forcing the control policy towards the default. This is the standard KL-regularized RL objective [Tishby and Polani, 2011, Rubin et al., 2012, Todorov, 2009, Fox et al., 2016, Levine, 2018] when using a learned default policy *π*_0_ conditioned on action history rather than a fixed unconditional marginal. This is an instance of conditional policy compression [Lai et al., 2025], which regularizes the policy toward a default conditioned on side information. Conditioning on the previous action lets the default store which action follows which, so it can chain actions into sequences without relying on external cues.

#### 2.1.2 Optimal policy and learning rules

##### Optimal value function

For state *s*, *V*^∗^ (*s*) is the expected total return (reward minus discounted KL cost) under the policy *π* that maximizes *J*:

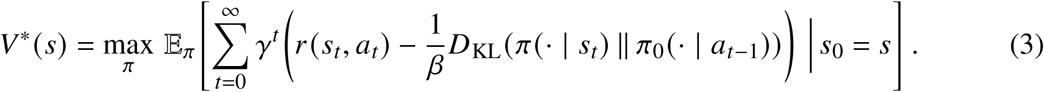

##### Bellman decomposition

Define the action-value

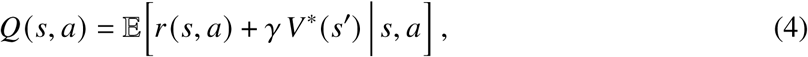

which absorbs all future expected reward from (*s*, *a*) onward. Expanding *D*_KL_ as a sum over actions, we arrive at the following optimization problem:

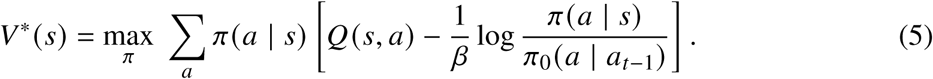

##### Pointwise solution

Forming the Lagrangian with multiplier *λ* for the normalization constraint Σ*_a_ π*(*a*|*s*) = 1 and differentiating with respect to *π*(*a* | *s*), we obtain:

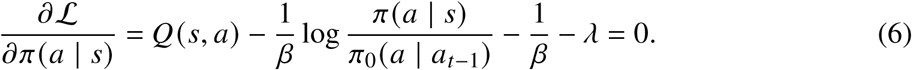

Solving for *π*(*a* | *s*) and absorbing the *a*-independent factor exp(−1 − *βλ*) into the normalization constant gives the optimal capacity-limited policy:

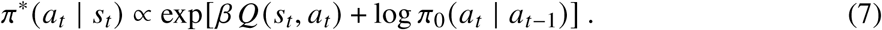

Eq. 7 carries an implicit unit coefficient with log *π*_0_: the value-based update is scaled by *β*, while the cached default policy enters at a fixed unit weight, independent of *β*. We make this coefficient an explicit parameter *τ*, so the policy takes the form

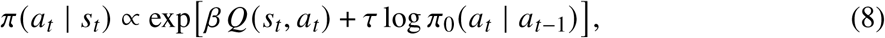

where *τ* = 1 recovers the normative policy derived (7).

Although the derivation recovers *τ* = 1, we retain it as an explicit parameter because it describes the influence of the default policy on the control policy. We model DLS damage as reducing the default’s weight (*τ* → 0), removing cached, history-dependent action selection, whereas DMS damage increases the default’s influence, leaving behavior dominated by the cached default when reward-driven learning is impaired. Empirical fits of policy compression models to behavior also show *τ* scattered around unity across subjects, with 89–96% of estimates falling between 0.5 and 1.5 across datasets [Gershman, 2020], motivating its treatment as a free parameter.

To compute *V*^∗^ (*s*) explicitly, we substitute the soft normative policy (7) back into the Bellman pointwise equation (5):

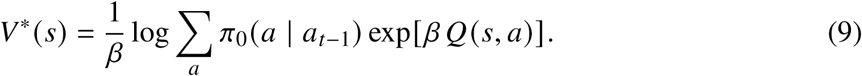

By definition, *Q*(*s*, *a*) is the expected immediate reward plus discounted future value:

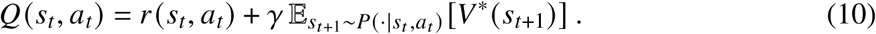

With *V*^∗^ given in closed form by (9), this becomes computable: the right-hand side depends only on *r*, the next state *s_t_*_+1_, and the current *Q* estimates (through *V*^∗^).

Since the agent is model-free, the transition *P* is unknown. Instead, the agent samples (*s_t_*, *a_t_*, *r_t_*, *s_t_*_+1_) from the environment and updates *Q* via the temporal-difference (TD) error [Sutton and Barto, 2018]; with *V*^∗^ substituted from (9), this is the reward-form analog of the G-learning update [Fox et al., 2016]:

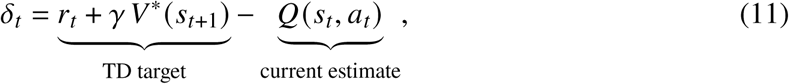

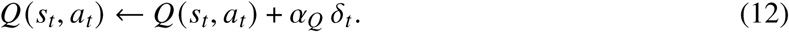

Note that *Q* is updated using only reward and the bootstrapped value *V*^∗^; the KL compression cost does not enter the *Q* update. This makes *Q* a cost-insensitive action value, following Fox et al. [2016]. Keeping *Q* cost-insensitive cleanly separates the reward-driven component (encoded by *Q*, learned in DMS) from the compression-driven component (encoded by *π*_0_, hypothesized in DLS), and is required for the boxed soft-policy form (7) to hold without modification.

##### Default policy update

Following Lai and Gershman [2021, eq. A.9], the default policy *π*_0_ is updated by a moving average towards the current control policy *π*, conditioned on the previous action:

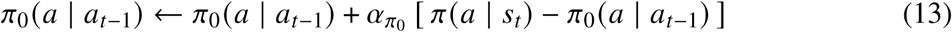

where *α_π_*_0_ ∈ (0, 1) is the default-policy learning rate. This causes *π*_0_(· | *a_t_*_−1_) to track the agent’s recent control-policy distribution following each *a_t_*_−1_, providing the compression target for *π*.

##### Balancing reward and complexity cost via *β* (tonic dopamine)

The parameter *β* plays the role of both a regularization coefficient for learning and an inverse temperature controlling exploration [Sutton and Barto, 2018, Rubin et al., 2012]. When *β* → ∞, the complexity cost is eliminated and the learning algorithm is equivalent to standard Q-learning. Empirically, this means that the agent’s actions are chosen solely based on external contingencies, not action history. When *β* = 0, the control policy is forced to be equal to the default policy, *π* = *π*_0_. In other words, the agent relies completely on prior action history to select future actions.

We interpret *β* as tonic striatal dopamine. Theoretical work proposes that tonic dopamine reports the average reward rate [Niv et al., 2007], and voltammetric and microdialysis measurements show that striatal dopamine levels track recent reward rate [Hamid et al., 2016, Mohebi et al., 2019]. Dopamine levels also set how strongly choices exploit learned values: elevating tonic dopamine increases exploitation of reward learning [Beeler et al., 2010]. In our framework, *β* scales the reward-driven value term and sets the balance between the two striatal streams: the value *Q*(*s*, *a*) (learned by DMS) and the default *π*_0_ (cached by DLS). As an agent learns a cued task, reward becomes frequent and predictable; tonic dopamine, and hence *β*, rises, so the policy is dominated by the value term. When reward becomes less predictable, as when external cues are removed after initial training, tonic dopamine falls, *β* decays, and control shifts towards the state-invariant default *τ* log *π*_0_.

In practice we set *β* close to 0 initially, when the *Q* estimates are still noisy, and increase it so the agent asymptotically obtains maximal reward [Fox et al., 2016], using a linear schedule

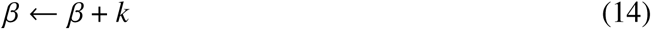

for constant *k* ≥ 0. In tasks where cues are removed after initial training (e.g., during automatization of motor sequences), we allow *k* < 0 so that *β* decays, implementing a reduction in tonic dopamine described above. This captures the empirical observation that automatization is accompanied by a transition from DMS- to DLS-dominated control [Turner et al., 2022], and reflects that tonic dopamine—and hence the agent’s willingness to encode state-specific information—decreases under reward uncertainty [Mikhael et al., 2021, Bari and Gershman, 2023].

We treat this linear schedule as a minimal placeholder for reward rate dependent changes in tonic dopamine rather than a mechanistic model. We also use a single *β* for both regions: *β* is a global gain on reward-driven value at the point where the two streams converge, not a DMS-specific dopamine level. A fixed-*β* control (*k* = 0) shows that no single constant *β* reproduces all four paradigms (Section 3.6).

### 2.2 Mapping the framework onto basal ganglia circuits

The basal ganglia has long been modeled as a central brain structure underlying reinforcement learning, with striatal circuits learning action values and arbitrating between flexible and habitual control [Joel et al., 2002, Daw et al., 2005, Frank, 2011, Bogacz and Gurney, 2007]. We ground this computational division in the anatomy of corticostriatal circuits, which are organized into parallel anatomical loops [Alexander et al., 1986, McGeorge and Faull, 1989, Hunnicutt et al., 2016]. The DMS (associative striatum) receives its cortical input from prefrontal cortex [McGeorge and Faull, 1989, Hintiryan et al., 2016], which encodes state and context [Miller and Cohen, 2001, Hampton et al., 2006], whereas the DLS (sensorimotor striatum) receives input from motor cortex [McGeorge and Faull, 1989, Hintiryan et al., 2016], which encodes actions and movements [Georgopoulos et al., 1986, Churchland et al., 2012](Figure 1).

Functionally, the DMS supports flexible, goal-directed behavior while the DLS supports habitual, history-dependent responding [Yin and Knowlton, 2006, Balleine et al., 2007, Graybiel, 2008, Graybiel and Grafton, 2015]. These two variables of our model mirror this anatomical distinction: the reward-driven value *Q*(*s*, *a*) is learned by DMS — it is conditioned on state and matches the state and context signal carried by PFC into DMS, while the default policy *π*_0_(*a* | *a_t_*_−1_) is encoded by DLS — conditioned on the previous action, matching the action signal carried by motor cortex into DLS (Figure 1). The balance between the two streams is set by the default-policy weight *τ* and the inverse temperature *β*, which we interpret as tonic striatal dopamine. We test this mapping in four canonical DLS–DMS dissociations: outcome devaluation [Yin et al., 2004], serial spatial reversal [Castañé et al., 2010], skilled action sequencing [Turner et al., 2022], and motor sequence execution [Mizes et al., 2023].

#### Lesion implementations

We propose that the soft policy (Eq. 7) is computed at a downstream site where the output of DMS and DLS converges — canonically at the thalamocortical targets of the basal ganglia output nuclei SNr/GPi [Mink, 1996, Aoki et al., 2019]. We interpret *τ* as the weight of the contribution of DLS (default policy) at the output node, while the contribution of DMS is scaled by the inverse temperature *β*; the balance between the two streams controls the selection of action. In the intact striatum (sham), the normative derivation places the DLS contribution at its optimal value *τ* ≈ 1 [Gershman, 2020], and lesions displace it away from unity.

We implement each lesion as a change in the affected region’s contribution to the policy. A DLS lesion removes the log *π*_0_ substrate from the readout, collapsing the weight of the default policy (*τ* → 0) and leaving the behavior purely value-driven *π*(*a*|*s*) ∝ exp[*βQ*(*s*, *a*)]. A DMS lesion impairs reward-driven value learning, modeled as a reduction in value-learning rate 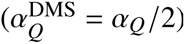; with the value signal weakened, action selection falls back on the preserved default, which we capture as a modest upward shift in the weight (*τ* slightly above 1). To summarize, we implement lesions in the following way across all four paradigms:

- **Sham**: Optimal policy in effect (Eq. 7); the normative optimum is *τ* = 1, and each agent draws *τ* ∼ N (1, 0.05).
- **DLS lesion**: Reduce *τ* to down-weight the contribution of log *π*_0_. When *τ* → 0, the policy collapses to *π* ∝ exp(*βQ*), thereby removing the influence of the default policy on behavior.
- **DMS lesion**: Reducing the rate of action-value learning (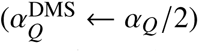), reflecting empirical evidence that DMS lesions impair the updating of state-action values. The loss of DMS counterweight at the SNr/GPi output node [Turner et al., 2022] causes DLS to dominate action selection (implemented by effective *τ* > 1).

When the lesion is applied, *τ* is drawn from the lesion-specific distribution and held fixed for the remainder of the simulation; Sham and pre-lesion agents draw *τ* ∼ N (1, 0.05). DLS lesions draw *τ* ∼ N (0, 0.05), effectively removing *π*_0_ from the policy; DMS lesions draw *τ* ∼ N (1.05, 0.05) and additionally slow value learning (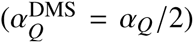). We use the same parameter distributions across all four paradigms. The timing of the lesion follows that of each experiment, as specified in the following paradigm sections.

### 2.3 Model parameters

All simulations share the same learning and lesion parameters (Table 2); paradigms differ only in task structure and training durations. The robustness of each dissociation to the lesion parameters (*τ*, *α_Q_*) is shown in Figure S1.

**Table 1:**
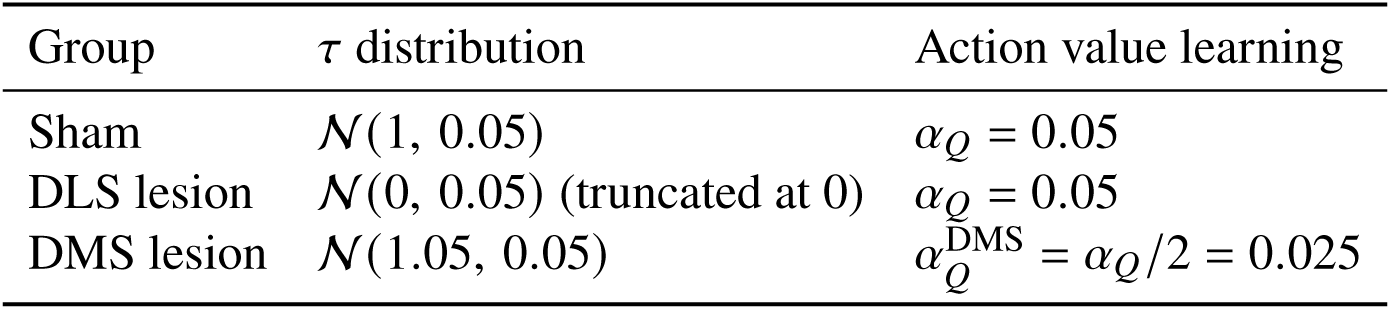
Lesion implementations. Each agent draws *τ* once from its group’s distribution at the time of lesion and holds it fixed thereafter.

| Group | *τ* distribution | Action value learning |
|---|---|---|
| Sham | 𝒩(1, 0.05) | *α*<sub>*Q*</sub> = 0.05 |
| DLS lesion | 𝒩(0, 0.05) (truncated at 0) | *α*<sub>*Q*</sub> = 0.05 |
| DMS lesion | 𝒩(1.05, 0.05) | *α*<sub>*Q*</sub><sup>DMS</sup> = *α*<sub>*Q*</sub>/2 = 0.025 |

**Table 2:**
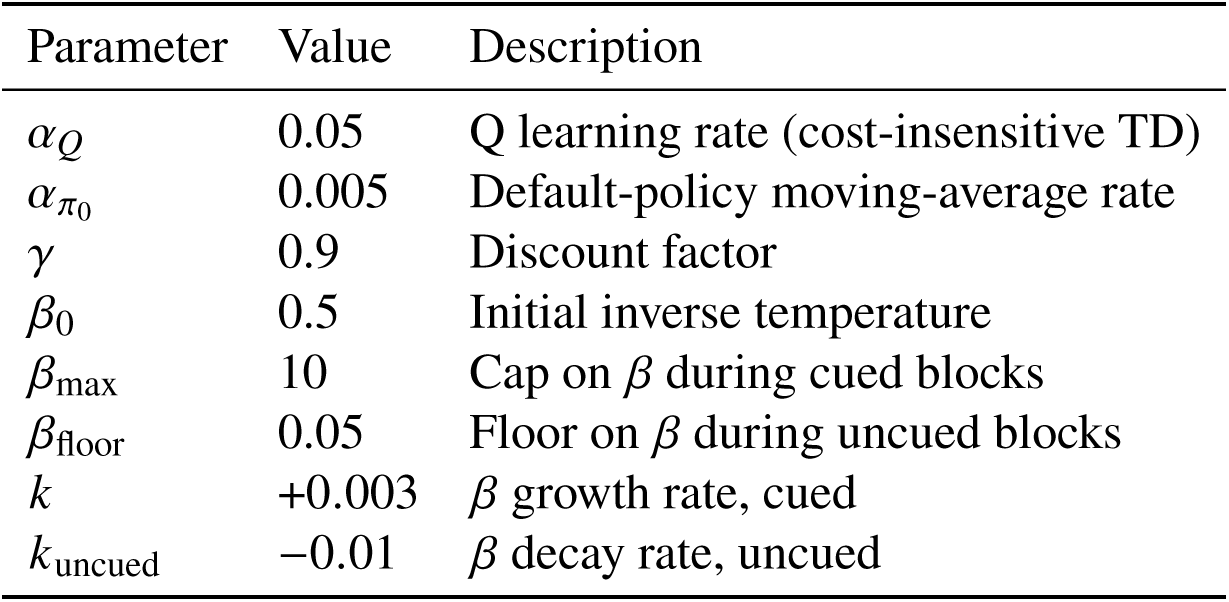
Model parameters, shared across all four behavioral paradigms.

| Parameter | Value | Description |
|---|---|---|
| *α*<sub>*Q*</sub> | 0.05 | Q learning rate (cost-insensitive TD) |
| *α*<sub>*π*<sub>0</sub></sub> | 0.005 | Default-policy moving-average rate |
| *γ* | 0.9 | Discount factor |
| *β*<sub>0</sub> | 0.5 | Initial inverse temperature |
| *β*<sub>max</sub> | 10 | Cap on *β* during cued blocks |
| *β*<sub>floor</sub> | 0.05 | Floor on *β* during uncued blocks |
| *k* | +0.003 | *β* growth rate, cued |
| *k*<sub>uncued</sub> | −0.01 | *β* decay rate, uncued |

### 2.4 Behavioral paradigms

All four simulations use the same policy (Eq. 8), with *τ* and *α_Q_* sampled separately for each lesion group (Table 1), but held fixed across paradigms (Table 2). The paradigms differ only in their task structure and in the time at which the lesion is applied, such that each paradigm isolates a different facet of the striatal contribution to behavior. The training durations were chosen so that the intact agents reached asymptotic performance before each test or lesion, reflecting the extended training in the original studies; they were not adjusted per lesion group.

#### 2.4.1 Outcome devaluation [Yin et al., 2004]

In outcome devaluation [Yin et al., 2004], a well-learned lever-press is challenged by devaluing its outcome, testing whether behavior remains flexible and goal-directed — declining after devaluation — or has become habitual and persists regardless. Following Niv et al. [2006], we treat motivational state as part of the internal MDP. The state is *s* = (*s*_ext_, *m*), where *s*_ext_ = 1 is external state and remains constant throughout the experiment because the lever is always available and the internal motivational state is *m* ∈ {*m*_1_, *m*_2_}: *m*_1_ is valued (sucrose desirable) and *m*_2_ is devalued (sucrose paired with LiCl induced malaise). Therefore, all state-dependence in this task comes through *m*. The action space is A = {press, no-press}, and the reward is provided only for pressing in the valued state, *r* ((1, *m*_1_), press) = 1, with all other pairs (*s*, *a*) producing zero. Following Yin et al. [2004], lesions are applied before training; agents acquire the task already lesioned. The experiment unfolds in three phases:

##### Training

Agents experience *s*_ext_ = 1, *m* = *m*_1_. *Q*(·, *m*_1_, press) rises and *π*_0_(press | press) consolidates toward 1; because *m*_2_ is never visited, *Q*(·, *m*_2_, ·) remains at its zero initialization throughout training. Training consists of 40 sessions of 300 trials.

##### Outcome conditioning

Yin et al. [2004] paired sucrose with LiCl for the devalued cohort between training and test; we mirror this by shifting that cohort’s internal motivational state to *m*_2_. The valued cohort stays in *m*_1_. The external state *s*_ext_ = 1 is unchanged.

##### Extinction test

All learning is frozen and the agent selects actions with no reward delivered. The valued cohort tests in *m*_1_, where high *Q*(·, *m*_1_, press) and consolidated *π*_0_ from training stage both bias toward pressing. The devalued cohort tests in *m*_2_; because training was with sucrose, *m*_1_, and agents never visited *m*_2_, the corresponding *Q* row remains at its zero initialization, and behavior in the test is therefore driven by *π*_0_ alone (or, for DLS-lesioned agents, by a uniform softmax over those zero *Q* values). Each test consists of 150 trials. We report test press rate as a percentage of each animal’s training-phase rate, matching the normalization used by Yin et al. [2004] (*n* = 25 simulated animals per group, mean ± SEM).

#### 2.4.2 Serial Spatial Reversal [Castañé et al., 2010]

In serial spatial reversal [Castañé et al., 2010], the rewarded lever repeatedly flips, requiring flexible relearning, which competes against perseveration on the cached default policy.

We model the task as a single-state, three-action MDP with S = {*s*_1_} levers available, A = {left, right, no press}, and reward *r* (*s*_1_, *a*^★^) = 1 for the currently rewarded lever (zero otherwise). With a single state, *Q* collapses to a length-three vector and *π*_0_(*a* | *a_t_*_−1_) to a 3 × 3 transition matrix. After the post-surgery retention phase, *α_π_*_0_ is halved to reflect that once a behavior is consolidated, the default policy updates are slower. Similar to Castañé et al. [2010], all simulated animals have identical training before they undergo lesion manipulations.

##### Training

An intact agent learns with *a*^★^ = left. Sessions consist of 70 trials, scored in 10-trial blocks, and the agent runs to a criterion of 9/10 correct in a block (or up to 50 sessions). The trained (*Q*, *π*_0_, *β*) at criterion is saved and used as the starting state for all three lesion branches.

##### Surgery and retention

The lesion is applied after the training phase and a retention phase reruns the same contingency to criterion to confirm post-surgery retention of the behavior. Retention runs for up to 5 sessions.

##### Serial reversals

Following post-surgery retention on the original contingency, the rewarded lever flips left ↔ right at the start of each of three serial reversals. On each reversal the agent runs to criterion (9/10 correct presses in a 10-trial block; up to 50 sessions), followed by a retention session on the new contingency before the next reversal begins. Following Castañé et al. [2010], each session was classified by a two-tailed binomial test against chance: errors in sessions with a higher chance response to the previously rewarded lever (≥ 44/70 presses) were counted as perseverative, errors in sessions at chance as learning errors, and sessions below chance were considered accurate.

#### 2.4.3 Skilled action sequencing [Turner et al., 2022]

In skilled action sequencing [Turner et al., 2022], a multi-step sequence becomes automatic once cues are removed, testing whether acquisition of automatic execution depends on the cached default rather than on reward-driven value learning. We model the task as a 5-state, 5-action MDP with S = {1, 2, 3, 4, 5} corresponding to step cues (latent during uncued phases) and A = {1, 2, 3, 4, 5} the five poke holes. Reward is delivered upon completing the full sequence (*r*_term_ = 1). To capture the gradual consolidation of motor habits across the early stages of training, *π*_0_ is initialized at the start of the uncued punished phase with partially consolidated chain transitions; this initialization is set identical across all lesion groups. As in Turner et al. [2022], lesions are applied before training: agents acquire the task after lesions and the study investigates how different striatal lesions impact the learning of automatised action sequence. The simulations unfold in three phases:

##### Cued training (Phase 1)

Each step *k* presents cue *s* = *k*, and the agent samples actions until it pokes the correct hole; wrong pokes incur *r* = 0 and update *Q* and *π*_0_ but do not abort the trial. *Q* learns cue→action mappings, while *π*_0_ tracks the policy via moving average, gradually forming the sequential chain. Phase 1 consists of 50 sessions of 100 trials.

##### Uncued sequencing (Phase 2)

Cues are removed and *s* becomes latent; the agent acts using a belief-state value 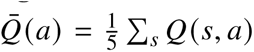. *Q* is frozen while *π*_0_ continues to update, and *β* decays under the annealing schedule. Errors do not abort the trial. Phase 2 consists of one session of 200 trials, serving as a brief transition from cued to uncued execution before phase 3, in which performance is measured.

##### Uncued punished (Phase 3)

Same belief-state structure as Phase 2, but any error aborts the trial immediately and only perfect 5-step completions earn reward. Phase 3 consists of 15 sessions of 200 trials, analyzed in five blocks of three sessions as in Turner et al. [2022].

#### 2.4.4 Cued vs Automatic motor sequence execution [Mizes et al., 2023]

In cued versus automatic motor sequencing [Mizes et al., 2023], the same motor elements are used to execute the sequence either by following cues or automatically from memory. Because the two modes share identical motor elements, the paradigm isolates the necessity of each striatal region for cued versus automatic execution.

We model the task as a 3-state, 3-action MDP with S = {*L*, *C*, *R*} (cue) and A = {*L*, *C*, *R*} (press). During uncued blocks, *s* is latent and the agent acts on the belief-state value under a uniform prior over the latent state, 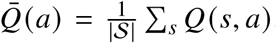. This belief-state *Q̅* is approximately uniform across actions once *Q̅* has been learned symmetrically, so action selection in uncued blocks is driven primarily by the default policy *π*_0_. The default policy *π*_0_(*a* | *a_t_*_−1_) is conditioned on the action history *a_t_*_−1_ (the previous press) and updated by a moving average toward the current control policy. Lesions are applied after training to study the necessity of striatal regions in execution of motor sequences.

##### Cued training

Stages 1–2 form a shared training trunk. Stage 1 is single cue–action association training with reward on pressing the cued lever, raising *Q* for the cued actions. Stage 2 is random 2-cue, 2-press training with reward dispensed upon pressing the two cued levers in a sequence, under which *Q* generalizes across cue order and *π*_0_ begins to track multi-step transitions. Agents then advance to a random 3-cue, 3-press stage, learning to follow the cues to produce three-press sequences with reward at sequence completion. Stage 1 lasts 20 sessions of 300 trials, Stage 2 20 sessions of 100 trials, and the random 3-cue stage 30 sessions of 100 trials. Because the cue order varies trial to trial, *π*_0_ does not cache a useful chain and the task is solvable only if the agent learns *Q*. We then apply the lesion and measure full-sequence accuracy on the random 3-cue task with learning frozen, isolating the necessity of each striatal term for cue-driven execution.

##### Automatic training

Agents are trained on a fixed 3-lever cued sequence with end-of-sequence reward, driving *π*_0_ to consolidate the sequential chain. Following the protocol of Mizes et al. [2023], the cues are then removed and agents perform the same sequence uncued, forcing them to rely primarily on *π*_0_. Cued training on the fixed sequence lasts 500 sessions of 100 trials, reflecting the extended overtraining of the automatic sequence in Mizes et al. [2023], followed by 30 uncued sessions of 100 trials. The uncued phase is shorter because, once cues are removed, *Q* is frozen (Section 2.5) and the sequence is already consolidated in *π*_0_; this phase serves as the transition to automatic execution. We then apply the lesion and measure full-sequence accuracy on the uncued task, with learning frozen, over 100 test trials.

### 2.5 Paradigm-specific overrides

After the initial training phase and post surgery retention of Castañé et al. [2010], *α_π_*_0_ is halved. This captures a basic property of consolidated behavior: once an action pattern has been well practiced and cached as default, it becomes harder to overwrite. This matters in Castañé et al. [2010] because its serial reversals repeatedly change the identity of the rewarded lever, and that relearning should be driven by a reward-based value update (*α_Q_*, DMS), which flexibly incorporates the new contingency, while the default (*π*_0_, DLS) resists change. We do not apply this override in the other paradigms because none of them invert a previously learned mapping. In Turner et al. [2022] and Mizes et al. [2023], training proceeds through progressive stages—involving a shift from per-press to sequence-completion reward, and a transition from cued to uncued execution—but none of these flips an already acquired reward contingency. Pressing the cued lever remains the right choice throughout; changing the reward schedule only defers the payoff, and uncued, automatic execution only consolidates the sequence rather than contradicting it. The default policy is therefore only ever built up on top of what was learned and never overwritten. Only in Castañé et al. [2010], where the rewarded lever repeatedly flips, is the cached default placed in direct conflict with the new reward contingency, and hence resists change.

In two of the task paradigms that we examine [Turner et al., 2022, Mizes et al., 2023], the external state *s* is latent: visual cues that would normally specify the appropriate action are removed, requiring the animal to execute an action sequence from memory. Hence, agents act according to a belief-state value computed under a uniform prior over latent states:

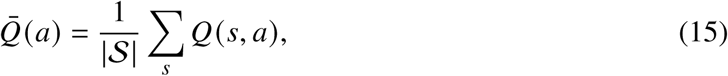

substituted for *Q*(*s*, *a*) in the soft policy (7). *Q* is frozen during uncued blocks (no observable *s* index for the TD update); *π*_0_ continues to update.

### 2.6 Comparison models

We compared the full model with two alternative models and one variant of our own, each simulated on all four paradigms with the same training protocols, lesion timing and number of agents (*n* = 25 per group).

i. Vanilla soft-Q learning: the default policy is removed for all groups (*τ* = 0), so that *π* ∝ exp(*βQ*) (Figure S2).
ii. Dual control model: the model of Daw et al. [2005], in which a model-based (tree-search) controller, mapped to DMS, and a model-free (cached) controller, mapped to DLS, each maintain posterior uncertainty about their value estimates, and on each step the less uncertain controller selects the action. We use the same parameter values as published. A lesion removes the corresponding controller from action selection. The model includes a forgetting factor: at each time step, both controllers’ estimates decay toward their priors, so that uncertainty never collapses and arbitration can keep tracking a changing environment. We used both the published value (0.98) and a slower one (0.999), since faster forgetting limits how much the model can learn over long training.

For outcome devaluation and serial reversal we followed the task structure of Daw et al. [2005]. In outcome devaluation [Yin et al., 2004], states are: lever available, reward delivered, and two terminal states (reward obtained or no reward); actions are lever press and magazine entry. Pressing the lever leads to the reward delivered state, and entering the magazine from there leads to the reward obtained state, which yields *r* = 1 during training; any other action ends the trial without reward. Following [Daw et al., 2005], devaluation sets the reward distribution to Beta(1, 15) (expected reward ≈ 0.06) for the devalued cohort, and leaves it unchanged for the valued cohort. The tree-search controller uses the new value immediately, whereas the cached controller updates only the terminal state, leaving its cached value of pressing unchanged. As in our model, the test is run in extinction with learning off. In serial reversal [Castañé et al., 2010], the states are the same, and the actions are left lever, right lever and magazine entry. Pressing the currently rewarded lever leads to reward delivered, and entering the magazine leads to reward obtained (*r* = 1); pressing the other lever or entering the magazine first ends the trial without reward. The rewarded lever switches at each reversal.

For the two sequence tasks, states during cued performance are the step cues (five in skilled action sequencing [Turner et al., 2022], three in motor sequence execution [Mizes et al., 2023]), and a correct response advances to the next cue, as in our model. Once cues are removed in automatic sequencing, what the agent can observe determines whether the model can execute the sequence at all, so we tested two state representations.

*Position known*: the state is the agent’s position within the trial (how many responses it has made so far). This follows the logic of Daw et al. [2005], whose states also mark progress through a trial (e.g., lever available, reward delivered, reward obtained). In their task, however, each state is signaled by an observable event, whereas in an uncued sequence nothing in the environment indicates the current step: the agent must keep track of its own past responses, a memory requirement the dual-controller model does not address. Giving the agent its position therefore assumes this memory is perfect. Because the sequence is fixed, each position always requires the same response, so values learned earlier in training carry over, and both controllers continue to learn throughout.

*Position unknown*: the agent receives the same information as our model. In an uncued sequence, nothing signals position within a trial, hence the current step is latent, values are frozen, and both controllers act on the belief-state value *Q̅* (the average over steps). In our model, the missing history is instead carried by the default policy *π*_0_(*a* | *a_t_*_−1_), which conditions on the previous action. The position-known version thus gives the dual-controller model more information than our model receives and is the stronger test of it (Figures S3–S6).

(iii) Fixed *β* (variant of our model): to test whether the *β* schedule is necessary, we removed it (*k* = *k*_uncued_ = 0) and held *β* constant throughout training and test, at a low (1), intermediate (3) or high (10) value; all other parameters and lesion rules are unchanged (Figure S7).

### 2.7 Statistical Analysis

Each lesion group consists of *n* = 25 simulated agents, and figures show mean ± SEM across agents, matching the original studies. Robustness sweeps and model comparisons (Section 2.6) used the same number of agents per condition so that conditions differ only in the parameter being varied. Statistical tests were chosen to match each paradigm’s design. For outcome devaluation [Yin et al., 2004], we compared devalued versus valued group’s press rates within each lesion group. For serial spatial reversal [Castañé et al., 2010] and action sequencing [Turner et al., 2022], we compared each lesion group (DMS, DLS) against Sham on the paradigm’s error and sequencing metric respectively. All three paradigms used two-sample, two-sided *t*-tests (df = 48), with effect sizes reported as Cohen’s *d*. For cued-versus-automatic sequence execution [Mizes et al., 2023], we compared preversus post-lesion accuracy within animal, separately for cued and automatic sequences, using the two-sided Wilcoxon signed-rank test (*n* = 25). Significance was assessed at *α* = 0.05; exact test statistics, degrees of freedom, and *p*-values are reported in the Results and figure legends. All comparisons were planned and specified a priori, following the original experiments. All simulations were performed in MATLAB R2021a with random seed rng(2205) for reproducibility.

### 2.8 Data availability

All simulation and analysis code is available at https://github.com/cheshta2205/Striatum-Policy-Regularization. This study generated no new experimental data; empirical values shown for comparison were taken from the cited publications [Yin et al., 2004, Castañé et al., 2010, Turner et al., 2022, Mizes et al., 2023].

## 3 Results

### 3.1 Testing the framework across four behavioral paradigms

To test whether our policy regularization model reproduces canonical DLS–DMS dissociations, we simulated four behavioral paradigms: outcome devaluation [Yin et al., 2004], serial spatial reversal [Castañé et al., 2010], skilled action sequencing [Turner et al., 2022], and cued-versus-automatic motor sequence execution [Mizes et al., 2023]. We chose these paradigms because each isolates a distinct aspect of striatal function and highlights different contributions of striatum to learning and action selection. We simulate all four paradigms with one set of shared parameters (Table 2), and the same update and lesion rules (Eqs. 12,13).

### 3.2 Outcome devaluation

Outcome devaluation paradigms provide a canonical test of the balance between goal-directed and habitual control. One hallmark study [Yin et al., 2004] shows that goal-directed behavior is sensitive to changes in outcome value, whereas habitual behavior persists despite devaluation. In the context of policy regularization, this paradigm provides a natural setting to dissociate the contributions of the default policy and flexible value-based control in action selection. Lesions to the DLS disrupt habitual responding, rendering performance sensitive to devaluation, whereas lesions to DMS impair goal-directed control, leading to persistent lever pressing despite changes in outcome value [Yin et al., 2004].

We model the task as a single external state with two motivational states — valued (*m*_1_) and devalued (*m*_2_) — and two actions — press and no-press (Figure 2A). After training, the intact (Sham) agent’s value function concentrates towards pressing in the valued state and remains at zero in the never-visited devalued state (Figure 2B), while the default policy consolidates the press→press transition to ≈ 1 (Figure 2C).

**Figure 2:**
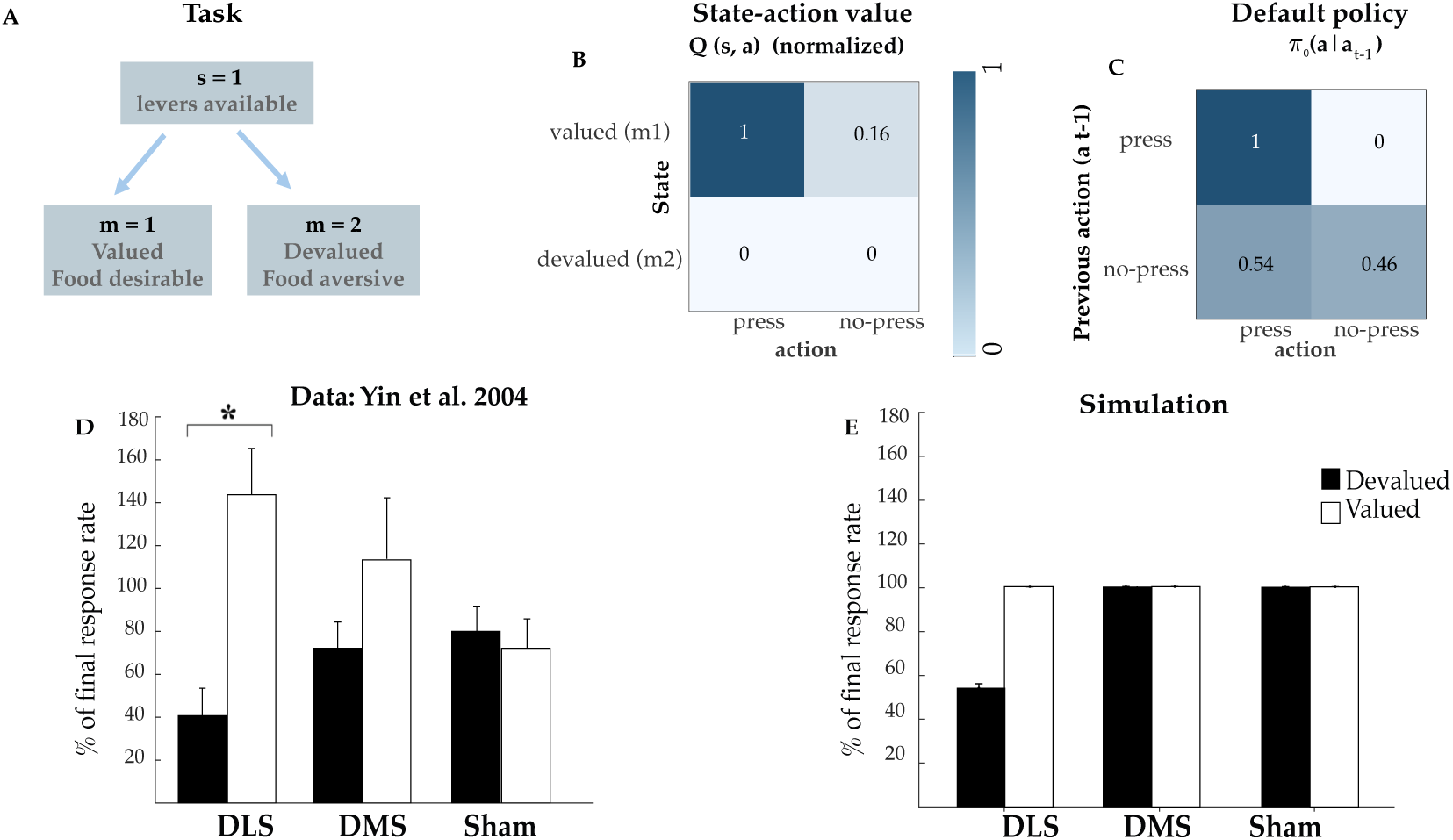
Outcome devaluation reproduces the canonical DLS–DMS dissociation. **(A)** Task: one external state (levers available) and two motivational states, valued (*m*_1_) and devalued (*m*_2_); actions are press and no-press. **(B)** Learned *Q*(*s*, *a*) (Sham, end of training; normalized): pressing has higher Q in valued state (*m*_1_) and zero in the never-visited devalued state (*m*_2_). **(C)** Learned default policy *π*_0_(*a* | *a_t_*_−1_): the press→press transition consolidates to 1. **(D)** Empirical data from Yin et al. [2004]: lever-press rate during extinction test, for valued and devalued cohorts across DLS-lesioned, DMS-lesioned, and Sham animals. **(E)** Simulation results under our policy regularization framework (*n* = 25 simulated agents per group, mean ± SEM; devalued vs. valued compared by two-sample two-sided *t*-test). Black: devalued cohort (tested in *m*_2_); white: valued cohort (tested in *m*_1_). Sham and DMS-lesioned agents are insensitive to devaluation (habit); DLS-lesioned agents reduce pressing in the devalued state (goal-directed).

Figure 2 D, E show that our model reproduces the canonical DLS–DMS dissociation: Sham and DMS-lesioned agents were insensitive to devaluation (devalued vs. valued press rate, two-sample two-sided *t*-test; Sham: 100.4 ± 0.02% valued vs. 100.4 ± 0.02% devalued, *t* (48) = −0.97, *p* = 0.34, *d* = −0.27; DMS: 100.5 ± 0.01% vs. 100.6 ± 0.02%, *t* (48) = 1.40, *p* = 0.17, *d* = 0.39), reflecting habitual lever pressing because of a preserved default policy. In contrast, DLS-lesioned agents were sensitive to devaluation, pressing at training-level rates when the outcome was valued but pressing less when devalued (100.5 ± 0.03% valued vs. 54.3 ± 1.9% devalued; *t* (48) = −24.65, *p* < 0.001, *d* = −6.97). With the default intact (*τ* ≈ 1), Sham and DMS agents press via the consolidated habit *π*_0_(press | press) = 1, which persists regardless of the outcome value; removing the default (*τ* → 0) leaves behavior governed by *Q*, and because the devalued state was never visited during training (*Q*(·, *m*_2_, press) = 0), pressing collapses. This dissociation is robust to lesion parameters — the habitual phenotype (as observed in DMS lesion/Sham) is invariant across the swept parameter range (*τ*, *α_Q_*), while the goal-directed behavior (as observed in DLS lesion) degrades smoothly with a rise in *τ* — suggesting that a partial DLS lesion can still allow habitual responding to develop (Figure S1A).

### 3.3 Serial spatial reversal

Reversal-learning paradigms test an agent’s ability to update a previously learned action policy when the reward contingency changes. Castañé et al. [2010] trained rats on a two-lever spatial discrimination task to learn that one lever was rewarded, applied striatal lesions (separate cohorts of DLS, DMS or sham lesions) and then tested animals across three serial reversals in which the rewarded lever identity switched. DMS-lesioned animals made substantially more perseverative errors, continuing to press the previously rewarded lever after reversal, whereas DLS-lesioned and Sham animals did not show this deficit.

In the context of policy regularization, this paradigm dissociates the contributions of the cached default policy *π*_0_ from the reward-driven action value *Q*. Because *Q* is updated from the reward feedback, it should be necessary to track the new contingency after the reversal. Thus, slowing *Q* updates in DMS-lesioned agents should produce perseverative errors. We model it as a single state with two levers whose rewarded identity flips across reversals (Figure 3A); the sham agent learns value concentrated towards the rewarded lever (Figure 3B) and a default policy consolidates towards it (Figure 3C). In contrast, DLS-lesioned agents retain intact reward-driven updating of *Q*, so reversal learning should remain intact. Removing *π*_0_ eliminates the influence of the previously cached default policy; therefore, the model predicts that DLS lesions should not increase perseveration.

**Figure 3:**
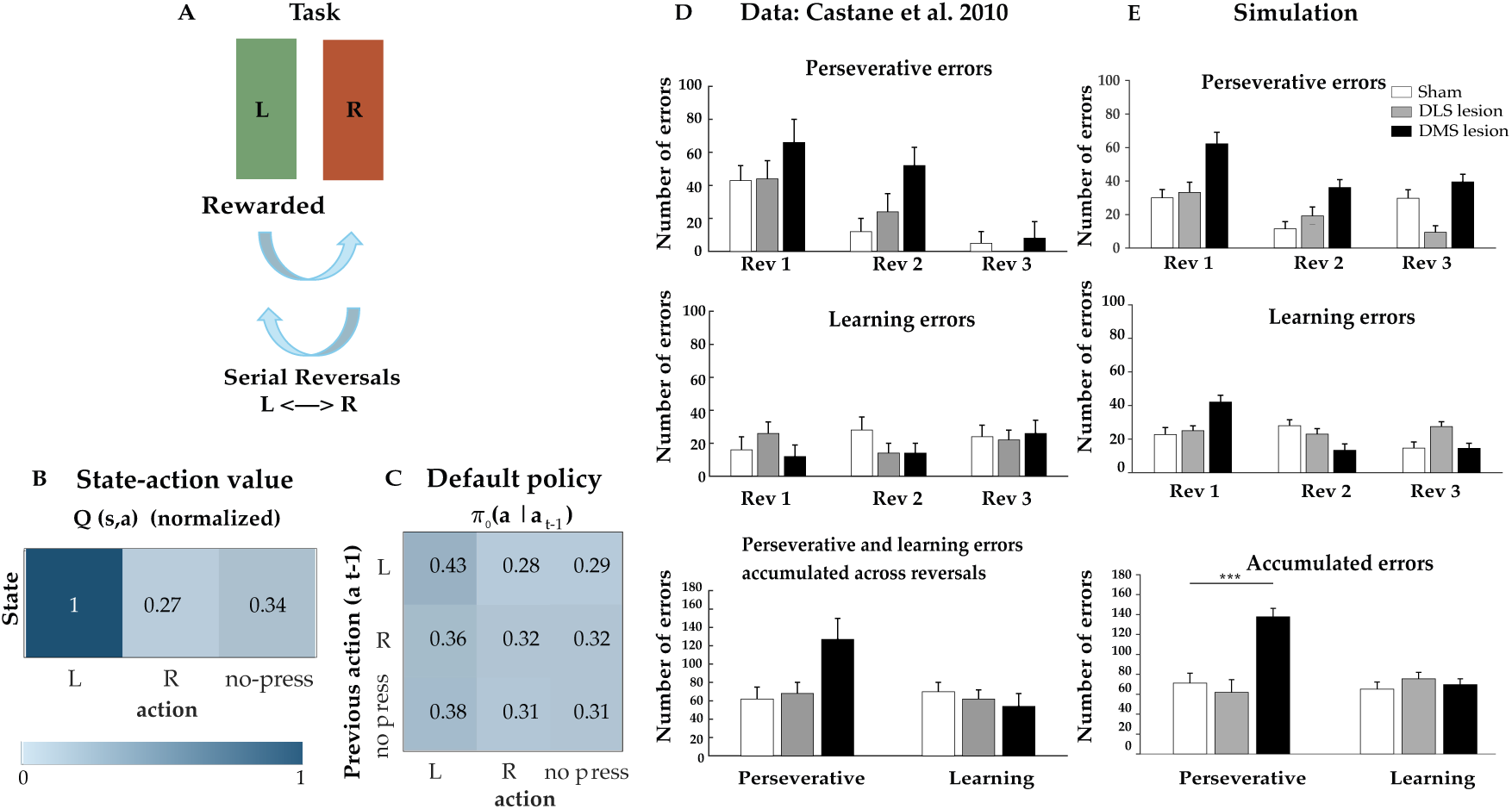
Serial spatial reversal reproduces the DMS-specific perseveration pattern of Castañé et al. [2010]. **(A)** Task: two levers (L/R) — the rewarded lever identity flips L↔R across serial reversals. **(B)** Learned *Q*(*s*, *a*) (Sham, end of Phase 1; normalized) — value concentrates towards the rewarded lever (L). **(C)** Learned default policy *π*_0_(*a* | *a_t_*_−1_): consolidated toward the trained lever, near-uniform elsewhere. **(D)** Empirical data: perseverative and learning errors across three reversals, for sham, DMS-lesioned, and DLS-lesioned animals. DMS lesions made significantly more perseverative errors. **(E)** Simulation results under our policy regularization framework (*n* = 25 simulated agents per group, mean ± SEM). Slowing *Q* learning (DMS) selectively elevates perseverative errors.

Our model reproduced this pattern (Figure 3D, E): DMS-lesioned agents perseverated far more than sham, while DLS-lesioned agents were indistinguishable from sham. Per reversal, DMS-lesioned agents made 45.9 perseverative errors, compared to 23.7 by Sham and 20.6 by DLS. DMS agents were impaired relative to Sham (two-sample two-sided *t*-test; *t* (48) = 5.13, *p* = 5.1 × 10^−6^, *d* = 1.45), whereas DLS agents did not differ from Sham (*t* (48) = −0.58, *p* = 0.56, *d* = −0.16).

This deficit was specific: learning errors did not differ across groups (Sham = 21.7, DLS = 25.1, DMS = 23.2; DMS vs. Sham *t* (48) = 0.49, *p* = 0.62, *d* = 0.14; DLS vs. Sham *t* (48) = 1.06, *p* = 0.30, *d* = 0.30), and the effect was largest on the first reversal (DMS = 62.2 vs. Sham = 29.9).

The model thus reproduces the central dissociation reported by Castañé et al. [2010]: DMS lesions selectively elevate perseveration, whereas DLS lesions spare reversal performance.

This deficit is driven by the value-learning rate: perseveration rises monotonically with a reduction in *α_Q_* (or slowing of value learning) but is invariant to *τ*, and raising *τ* alone (with normal *α_Q_*) leaves perseveration at Sham level (Figure S1B).

### 3.4 Skilled action sequencing

Action-sequencing paradigms probe how animals learn to stitch individual actions into coherent sequences. Turner et al. [2022] investigated the contributions of striatal regions in acquisition of automatic sequences in rats, using a five-step heterogeneous nose-poke task. Animals first learned the action sequence by following visual cues, then performed the same sequence without cues, relying on memory-guided execution. In the final punishment stage, sequences remained uncued, but any error aborted the trial, so that reward was only delivered after perfectly executed sequences. This task revealed opposing contributions of DMS and DLS in acquisition of skilled automatised sequence: DMS-lesioned animals produced more correct sequences per block than Sham throughout acquisition, while DLS-lesioned animals were impaired — producing a DMS > Sham > DLS ordering in correct sequences per block. In the policy-regularization framework, this task dissociates cached, history-dependent action chains stored in *π*_0_ from flexible, externally-guided control based on *Q*. Once external cues are removed, successful performance of the sequence depends on the consolidated action chain stored in *π*_0_. Agents that rely more strongly on this default policy should execute the uncued sequence more efficiently, whereas agents whose default is ablated (*τ* → 0) should be impaired.

The five-step sequence is illustrated in (Figure 4A). When entering the uncued stage, the sham agent has learned the association between cues and actions (Figure 4B) and has partially cached the sequential chain in *π*_0_ (Figure 4C).

**Figure 4:**
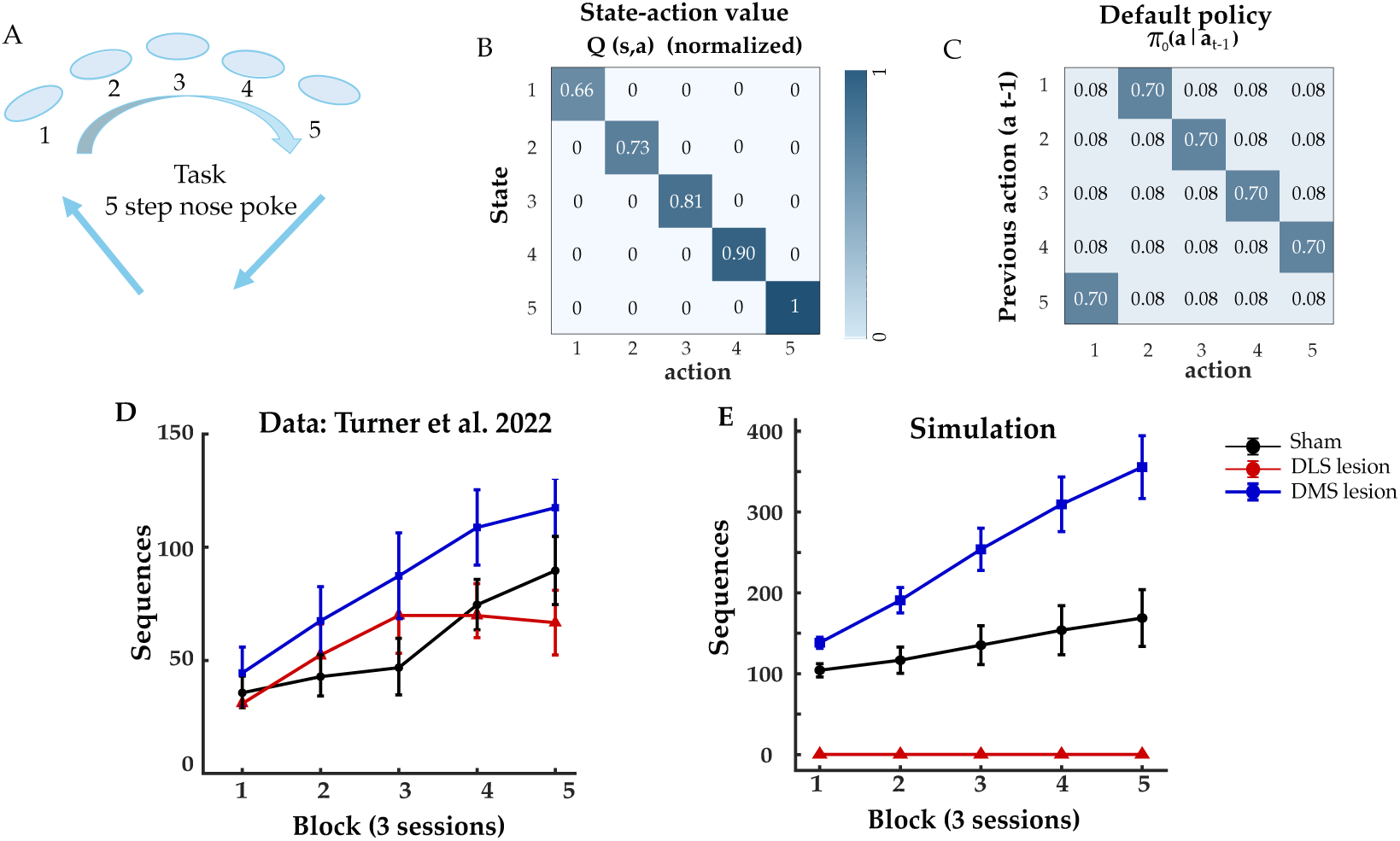
Skilled action sequencing reproduces the DMS > Sham > DLS ordering of Turner et al. [2022]. **(A)** Task: five-step nose-poke sequence (1→2→3→4→5). **(B)** Learned *Q*(*s*, *a*) (Sham, end of cued Phase 1; normalized): a clean cue→action diagonal – agent learns to follow external state (cues) to select actions. **(C)** Default policy *π*_0_(*a* | *a_t_*_−1_) entering the uncued test: a partially consolidated sequential chain, initialized identically for all lesion groups (Methods). The simulation therefore models how an already-formed chain is refined under uncued sequencing, with lesions acting on how strongly that chain is read out. **(D)** Empirical data: correct sequences per block during punished acquisition. **(E)** Simulation under the policy-regularization framework (*n* = 25 agents per group, mean ± SEM). DLS-lesioned agents (*τ* ≈ 0) are impaired because the cached *π*_0_ chain is ablated and *Q* is uninformative under uncued belief-state averaging; both Sham and DMS retain the action sequence and perform far above DLS. DMS-lesioned agents (*τ* ↑) lean more strongly on the consolidated chain and show the best performance. Partial DLS lesions are shown in Figure S8.

Our model reproduced the DMS > Sham > DLS ordering (Figure 4D, E). This pattern arises because once visual cues are removed, the belief-state value no longer carries information about the current action in the sequence. Sequence execution must therefore be entirely driven by the action chain stored in *π*_0_. DMS-lesioned agents, which rely more strongly on this cached policy, commit to the learned chain and show the best performance in both the first block (DMS = 138.3 vs. Sham = 104.3; *t* (48) = 3.19, *p* = 0.0025, *d* = 0.90) and the last block (DMS = 355.6 vs. Sham = 168.8; *t* (48) = 3.57, *p* = 8.2 × 10^−4^, *d* = 1.01).

In contrast, DLS-lesioned agents lack the contribution of the default policy (*τ* ≈ 0), leaving action selection dependent on a near-uniform softmax over *Q̅* and yielding near-chance step transitions; these agents produced almost no correct sequences during acquisition and performed significantly worse than Sham agents in both the first block (Sham = 104.3 vs. DLS = 0.2; *t* (48) = −12.73, *p* = 5.3 × 10^−17^, *d* = −3.60) and the last block (Sham = 168.8 vs. DLS = 0.2; *t* (48) = −4.81, *p* = 1.5 × 10^−5^, *d* = −1.36). Thus, our model recapitulates the results reported by Turner et al. [2022].

This ordering is robust to parameter sweep across a range: the DLS impairment persists across the entire swept *τ* range (Figure S1C). The model’s DLS lesion removes the default entirely (*τ* → 0) and so marks the upper bound of the deficit. To relate it to the residual performance of lesioned animals, we extended the sweep to partial DLS lesions (Figure S8). Five-step performance depends steeply and nonlinearly on *τ*: because the default supplies every transition, a small reduction in its weight is multiplied across the five steps, so performance stays low over most of the range and recovers sharply as *τ* approaches 1, reaching a level comparable to lesioned animals. The same reduction spares the three-step sequence of Mizes et al. [2023] (Figure S1D). The model therefore predicts that the effect of partial DLS lesions should depend on sequence length, with longer sequences disproportionately impaired.

The lesion magnitude at which behavioral recovery emerges can be mapped directly with graded DLS inactivation (titrated muscimol or optogenetic inhibition) and should be tested in future work. Notably, whereas *α_Q_* is critical for reversal, *τ* is the dominant parameter here, so each component of the DMS lesion is required for a different paradigm and neither can be dropped.

### 3.5 Cued vs. automatic motor sequence execution

Mizes et al. [2023] studied the contribution of striatal regions in execution of motor sequences. In particular, rats were trained to press three levers either in a randomly cued sequence (CUE) or in a predetermined, uncued sequence from memory (AUTO; Figure 5A). After learning both cued and automatic sequences, DMS or DLS was lesioned in separate cohorts of rats to test the necessity of each region in cued versus automatic motor sequence execution. They found that DLS lesions selectively impaired automatic sequence execution (AUTO) while sparing cued performance (CUE), whereas DMS lesions had no significant effect on either cued or automatic sequence execution.

**Figure 5:**
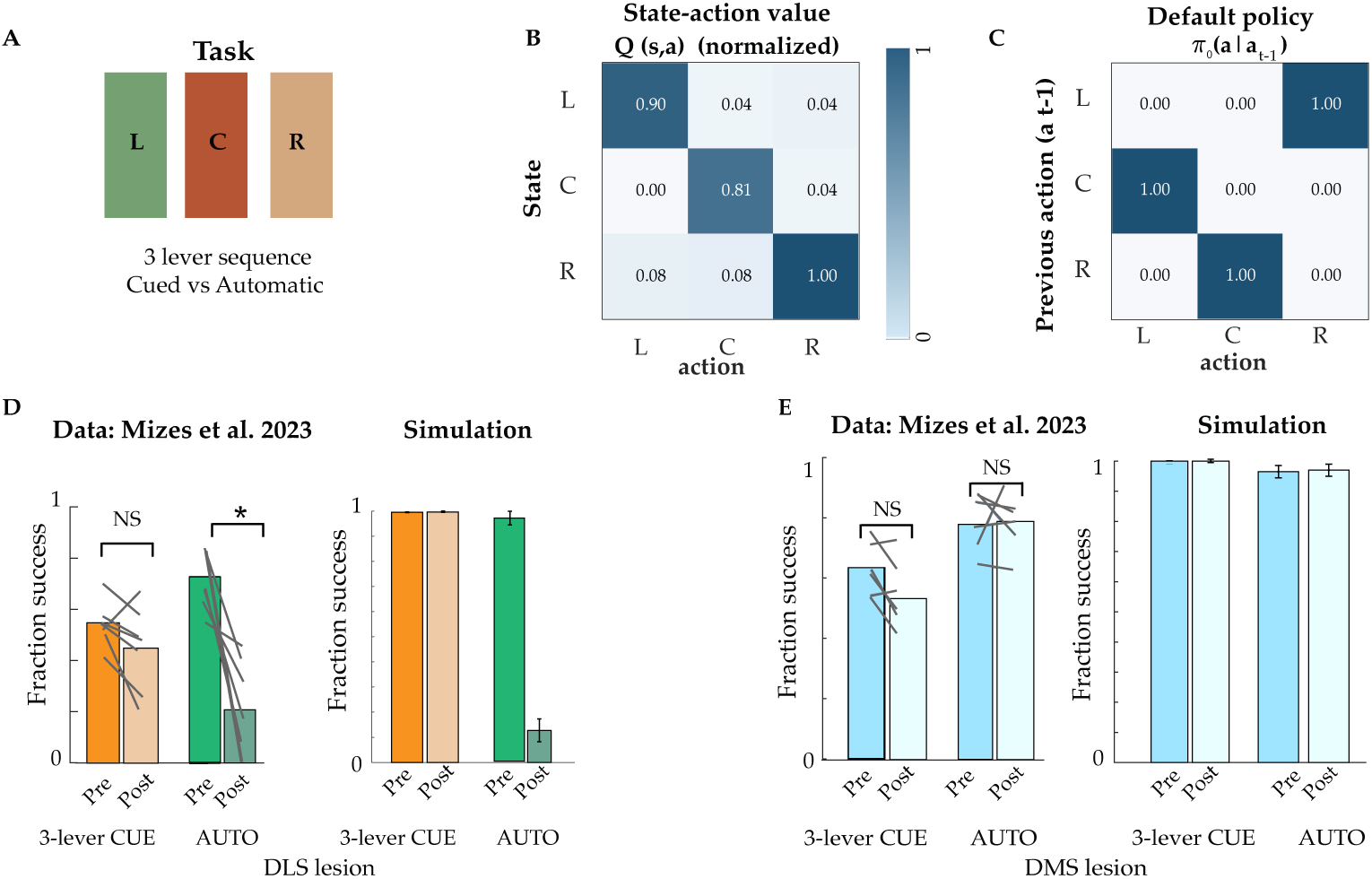
Cued vs. automatic execution reproduces the dissociation of Mizes et al. [2023]. **(A)** Task: three levers (L, C, and R); cued (random flexible sequences) vs. automatic (fixed uncued) sequence. **(B)** Learned *Q*(*s*, *a*) (intact, end of training; normalized): a cue→press diagonal: learned cue action mapping. **(C)** Learned default policy *π*_0_(*a* | *a_t_*_−1_): consolidated fixed 3-lever sequential chain. **(D)** Each panel plots full-sequence accuracy (pre- vs. post-lesion) on execution of randomly cued sequencing task and the uncued automatic sequencing task, with the empirical data of Mizes et al. [2023] (left) beside the matched simulation (right; *n* = 25 agents per group). DLS lesion (*τ* ≈ 0): the cued task is spared while the automatic task is significantly impaired, because removing the consolidated chain in *π*_0_ eliminates the only signal driving the uncued sequence. **(E)** DMS lesion (*α_Q_* ↓, *τ*↑): both tasks are spared, because the lesion is applied after *Q* has learned, so slowed value learning has no behavioral consequence.

In the context of policy regularization, this paradigm dissociates the two terms of the soft policy: Cued execution depends on learned *Q*(*s*, *a*) values, because trial-specific visual cues indicate what sequence of actions should be selected and automatic execution depends on a consolidated default policy *π*_0_(*a* | *a_t_*_−1_), because the animal must produce the sequence from action history after visual cues are removed. The intact agent learns cue-action associations required to perform cued sequences in cue→press (Figure 5B) and consolidates the fixed AUTO sequence in *π*_0_ (Figure 5C) as a cached pattern. Because lesions are applied after training, the model predicts that DLS lesions should selectively impair automatic execution by removing the stored default policy action chain (*τ* → 0), whereas DMS lesions should spare both tasks: cued execution because the cue-action *Q* values were learned before the lesion, which only slows further *Q* learning, and automatic execution because it relies on *π*_0_, which the DMS lesion leaves intact.

Figure 5D, E reproduces this selective DLS impairment. On the random 3-cue task, both groups maintained high performance after lesion (mean ± SEM full-sequence accuracy, pre- vs. post-lesion; Wilcoxon signed-rank, within-animal, *n* = 25; DLS: 0.995 ± 0.001 vs. 0.996 ± 0.001, *W* = 25, *p* = 0.46; DMS: 0.995 ± 0.001 vs. 0.998 ± 0.001, *W* = 28.5, *p* = 0.29). DLS lesions (*τ* → 0, removing *π*_0_) spared cued execution because *Q* alone follows the trial-specific cues; DMS lesions (*α*^DMS^ = *α_Q_*/2) caused no cued impairment because *Q* had already learned before the lesion. On the uncued automatic task, the two dissociated: DLS-lesioned agents had a significant drop (0.97 ± 0.03 vs. 0.13 ± 0.05; *W* = 325, *p* = 1.2 × 10^−5^), whereas DMS-lesioned agents were spared (0.96 ± 0.02 vs. 0.97 ± 0.02; *W* = 1.5, *p* = 0.19). Without cues, execution depends on the cached chain in *π*_0_; ablating it (*τ* → 0) collapses performance, while intact value learning is irrelevant once the task no longer relies on cue-driven *Q*. Thus, the model reproduces the key findings reported by Mizes et al. [2023]: execution of automatic motor sequence depends on DLS, whereas execution of cued sequences — which can be reconstructed trial-by-trial from sensory input — does not.

This dissociation is robust across a range of values for each parameter: DMS spares both cued and automatic execution across the full range of *τ* and *α_Q_* (flat ≈ 1), while DLS spares cued performance but impairs automatic execution at *τ* = 0, with performance progressively rescued as *τ* rises (Figure S1D). This graded rescue makes a testable prediction: partial DLS lesions should recover automatic execution of the three-step sequence, not the all-or-none loss of a complete lesion. Under a matched partial lesion, the model spares the shorter sequence of Mizes et al. [2023] but not the longer sequence of Turner et al. [2022] (Figure S1C,S8) — these paradigms differ in both sequence length and consolidation stage; we return to their interpretation in the Discussion.

### 3.6 Comparison with alternative models

We next asked whether the four dissociations require policy regularization or could arise from alternative accounts (Section 2.6).

#### Vanilla soft-Q learning

Removing the default policy abolished three of the four dissociations (Figure S2). In outcome devaluation, all three groups became devaluation-sensitive, so the habitual responding of Sham and DMS-lesioned agents disappeared (Figure S2A). In skilled action sequencing, no group executed the uncued sequence, and the DMS > Sham > DLS ordering was lost (Figure S2C). In cued versus automatic execution, cued sequences were performed normally, but automatic execution failed before and after the lesion in both groups (Figure S2D): without a cached chain, nothing drives the uncued sequence. Serial reversal was the exception. Vanilla RL still produced DMS-specific perseveration (Figure S2B), because the deficit reflects slow updating of *Q*, which vanilla RL retains. Without *π*_0_, however, intact and DLS-lesioned agents barely perseverated, whereas animals in all groups do [Castañé et al., 2010]. The default policy therefore accounts for the baseline perseveration of intact animals, not the DMS deficit itself.

#### Dual-controller model

The dual-controller model of Daw et al. [2005] reproduced outcome devaluation: removing the cached controller restored devaluation sensitivity, whereas removing tree search did not (Figures S3–S6). It did not reproduce the other three dissociations. In serial reversal, with the published forgetting factor no group perseverated (Figures S3, S5). With slower forgetting, DMS-lesioned agents perseverated more than Sham (20.2 vs. 10.8 errors per reversal, *p* = 0.002), but so did DLS-lesioned agents (17.5, *p* = 0.02), so the two lesions produced similar deficits rather than the DMS-specific perseveration seen in the data (Figures S4, S6). In both uncued sequence tasks [Turner et al., 2022, Mizes et al., 2023], when the agent’s position within the trial was unknown, as it is once cues are removed, no group executed the sequence (Figures S5, S6). When position was known, sequences were learned only with slower forgetting (0.999), not with the published value (0.98). Even then, DLS lesions spared both the five-step sequence (231 vs. 267 correct sequences in the last block, 87% of Sham) and automatic execution (0.78 vs. 0.77 pre- vs. post-lesion, *p* = 0.77), DMS lesions did not enhance five-step sequencing, and DMS lesions impaired cued execution (*p* = 0.003), unlike the data (Figure S4). This follows from the model’s structure: both controllers learn values over the same states, so either can execute a learned sequence on its own. After a DLS lesion, tree search can still plan the sequence from its learned model of the task, and no deficit appears.

#### Fixed *β* model

Finally, the *β* schedule was necessary: no fixed *β* reproduced all four paradigms (Figure S7). With low or intermediate *β* (1, 3), agents failed to perform cued sequences in the motor sequence task. With high *β* (10), five-step uncued sequencing collapsed in all groups, including Sham (last block ≈ 0), and perseveration after reversal was roughly sixfold higher than under the schedule.

## 4 Discussion

We have proposed that the DLS–DMS division of labor reflects a single computational principle: maximizing reward while minimizing the cost of deviating from a default policy. We formalized this trade-off using KL-regularized control, in which DMS supports reward-driven value learning via *Q*, while DLS stores a robust, history-dependent default policy *π*_0_ that caches previously consolidated action patterns. Under this framework, DLS “regularizes” DMS-driven control by biasing action selection toward cached behaviors that are efficient and robust, but less flexible when contingencies change.

A single model, with learning and lesion parameters shared across paradigms, reproduced canonical DLS–DMS dissociations across four paradigms: DLS lesions removed the default policy contribution, and DMS lesions slowed value learning and shifted control toward the default. Across these tasks, the model explains why DMS is necessary for flexible updating when reward contingencies change, whereas DLS is necessary for the development of cached structures. Table 3 summarizes the experimental dissociations reproduced by the model.

**Table 3:** Summary of simulated lesion results across the four paradigms, stated relative to Sham. In each case, the model reproduces the dissociation reported in the study.

| **Paradigm** | **DLS lesion (*τ* → 0)** | **DMS lesion (*α*<sub>*Q*</sub> ↓, *τ* ↑)** |
|---|---|---|
| Outcome devaluation [Yin et al., 2004] | Sensitive (goal-directed) | Insensitive (habit) |
| Serial spatial reversal [Castañé et al., 2010] | Unchanged | Elevated perseveration |
| Skilled action sequencing [Turner et al., 2022] | Impaired | Enhanced |
| Cued vs. automatic motor sequence execution [Mizes et al., 2023] | Cued spared, automatic impaired | Both spared |

Policy regularization provides a normative interpretation of skill automatization and habits: by caching action transitions that were once computed through reward-driven control, agents reduce the information cost of action selection and free cognitive resources for other demands [Haith and Krakauer, 2018, Du et al., 2022]. A key prediction of our framework is that when cognitive resources are limited, agents should be less able to maintain state-dependent policies and rely more strongly on default policies. This prediction is consistent with evidence that working memory load reduces instrumental learning rate (subserved by DMS in our model) and increases choice randomness [Park et al., 2023].

A related prediction, grounded in our simulations, is that reliance on the cached default policy—and hence on DLS—scales with the complexity of automatic behavior. *π*_0_ supplies the action at each step and degrading it corrupts each transition, so the deficit increases with sequence length, and behavioral recovery requires a milder lesion (Figure S1C, D, S8). The three-press task of Mizes et al. [2023] and the five-poke task of Turner et al. [2022] are ordered in this direction, though they also differ in learning stages tested, so sequence length and task complexity are not isolated. This is testable by combining graded DLS inactivation with sequences of varying length within a single paradigm, holding training history fixed.

Our model also formalizes the idea that DMS and DLS compete for behavioral control. Experimental studies have shown that these regions make opposing contributions to instrumental learning and habit formation [Yin et al., 2004, 2005], and DMS lesions enhance skilled sequencing while DLS lesions impair it [Turner et al., 2022]. This is consistent with neural recordings showing that DMS disengagement is predictive of habit formation [Thorn et al., 2010, Thorn and Graybiel, 2014, Kupferschmidt et al., 2017]. In our model, this competition is captured by the relative influence of *Q* and *π*_0_ on the policy. An interesting avenue for further exploration is how this balance changes over learning: *β* may fall and *τ* rise as behavior automatizes, mirroring the transfer of control from DMS to DLS. Here we use a simple linear schedule for *β* as a placeholder for reward-rate-dependent changes in tonic dopamine (Section 2.1); linking *β* to measured dopamine dynamics is an important next step.

### Why policy regularization rather than alternative models?

Comparing our model with vanilla soft-Q learning (*τ* = 0 throughout; Figure S2) shows that the default policy is necessary: the vanilla model failed to reproduce three of the four dissociations. Vanilla RL reproduced only the direction of the serial-reversal deficit, which rests on the reward-driven value term that standard RL contains, and even there intact and DLS-lesioned agents almost never perseverated, unlike animals. The dual-controller model of Daw et al. [2005] reproduced outcome devaluation but not the reversal or sequencing dissociations (Figures S3–S6). Because its two controllers learn values over the same states, either can execute a learned sequence alone, so neither lesion produces a selective deficit; and once cues are removed, it can execute a sequence only if its position in the sequence is supplied, which assumes a perfect memory of its own past responses. In our model, by contrast, the two terms condition on different information—*Q* on the external state, *π*_0_ on the previous action—so once cues are removed, only *π*_0_ carries the sequential structure. Unlike in the dual-controller model, where tree search can still plan the sequence after the cached controller is removed, no other component can take over after a DLS lesion, which makes DLS necessary for automatic execution. The *β* schedule was also necessary: no fixed *β* reproduced all four paradigms (Figure S7). Our model therefore predicts that tonic striatal dopamine should rise during cue-guided training and fall once cues are removed, and that manipulating it in either phase should disrupt the corresponding behavior. Parameters were shared rather than fitted per paradigm, and sweeping each lesion parameter preserves the qualitative pattern (Figure S1). The two DMS manipulations are each necessary: reduced *α_Q_* drives reversal perseveration, and increased *τ* drives enhanced automatic sequencing. Together, these comparisons suggest that policy regularization is a generalizable principle for the striatal division of labor.

### Relation to prior default-policy and compression models

Our model builds on a set of models that regularize a control policy towards a default policy. Piray and Daw [2021] shows that linear reinforcement learning, in which control cost is the KL divergence from the default policy, explains several aspects of grid fields in entorhinal cortex, habitual decision making, and cognitive control. The KL cost is closely related to the account developed here, though they did not apply this idea to explaining computation in the DLS and DMS. Moskovitz et al. [2024] derive a related two-policy architecture, showing that dual-process phenomena such as habitual action selection emerge from minimizing the description length of behavior. Their framework remains domain-general; the controller and default are not mapped to brain regions, and the default is not conditioned on action history. Our model grounds these two components in striatal regions.

### How do striatal circuits implement policy regularization?

One proposal is that transitions from DMS- to DLS-dominated control during habit formation are regulated by the “ascending spiral,” a striato-nigro-striatal circuit in which DMS activity influences dopamine signaling in DLS (i.e., DMS → SNr → SNc → DLS) [Haber et al., 2000, Yin and Knowlton, 2006, Lerner, 2020]. However, anatomical work suggests that this circuit may not support straightforward disinhibition of DLS dopamine signaling [Ambrosi and Lerner, 2022]. Instead, closed striatonigrostriatal loops may be better suited to support disinhibition. Other polysynaptic routes, including pathways through SNr, thalamus, and motor cortex, may also mediate communication between striatal subregions [Aoki et al., 2019]. Thus, while our model specifies the computational roles of DMS and DLS, the circuit mechanisms underlying their interaction remain an important target for future work.

### Neural signatures of a compressed policy

A central but untested implication of our model is that policy compression should be evident in striatal population activity. DMS activity should encode state-specific and reward-related information needed to support flexible *Q*-based learning. DLS activity, by contrast, should primarily encode previous actions and action-to-action transitions and carry less information about the external state. These signatures should also be dynamic: decoding current state and previous action from simultaneous DMS and DLS recordings during sequence learning should show DLS state dependence declining and action-history dependence increasing over training, while DMS remains state-dependent during flexible learning.

### Limitations

One limitation concerns the magnitude of the DLS lesion effect. Our model reproduces the ordering in Turner et al. [2022]—DMS > Sham > DLS—but produces a larger DLS effect than seen in lesioned animals. Because losses compound across the five steps, the model’s complete lesion lies at the severe end of a steep, nonlinear curve; partial reductions of the default bring performance into the range of lesioned animals (Figure S8), consistent with incomplete biological lesions and residual default control through other circuits. The gap also reflects the learning stage tested, since Turner et al. [2022] probe automaticity before the sequence is fully cached, and what the lesion disrupts: their deficit is in sequence initiation, whereas our model, acting from a state-averaged *Q*, treats the sequence as an all-or-none unit, so removing the default abolishes execution. The model also omits other regions—notably motor cortex, which initiates actions [Murakami et al., 2014] and is critical while a behavior is being learned [Kawai et al., 2015], and could sustain performance when caching is incomplete. Embedding policy regularization in a circuit-level model including these regions would predict DLS dependence emerging gradually with caching.

In conclusion, policy regularization provides a normative computational perspective for understanding how the brain balances flexibility and robustness in behavior. By mapping reward-driven control and learning onto DMS and compressed default policies onto DLS, the model explains why flexible learning and automatic action selection depend on partially competing striatal systems. This perspective links habit formation, skill automatization, and cognitive-resource constraints within a single formal account, and generates testable predictions about behavior and neural population codes.

## Acknowledgments

The authors thank Claude (Anthropic) for manuscript and code proofreading. All scientific ideas, analyses, and conclusions are the authors’ own.

## Supporting information

**S1 Appendix. Supplementary figures S1–S8.** Robustness of the lesion dissociations to lesion parameters (Figure S1); comparison with vanilla reinforcement learning (Figure S2); comparison with the dual-controller model of Daw et al. (2005) (Figures S3–S6); fixed-*β* simulations (Figure S7); and partial DLS lesions in skilled action sequencing (Figure S8).

## References

1. Abbas Abdolmaleki, Jost Tobias Springenberg, Yuval Tassa, Remi Munos, Nicolas Heess, and Martin Riedmiller. Maximum a posteriori policy optimisation. In International Conference on Learning Representations, 2018. URL https://openreview.net/forum?id=S1ANxQW0b.

2. Thomas Akam and Mark E Walton. What is dopamine doing in model-based reinforcement learning? Current Opinion in Behavioral Sciences, 38:74–82, 2021. ISSN 2352-1546. doi: 10.1016/j.cobeha.2020.10.010. URL https://www.sciencedirect.com/science/article/pii/S2352154620301558. Computational cognitive neuroscience.

3. G. E. Alexander, M. R. DeLong, and P. L. Strick. Parallel organization of functionally segregated circuits linking basal ganglia and cortex. Annual Review of Neuroscience, 9:357–381, 1986. doi: 10.1146/annurev.ne.09.030186.002041.

4. Priscilla Ambrosi and Talia N Lerner. Striatonigrostriatal circuit architecture for disinhibition of dopamine signaling. Cell Rep., 40(7):111228, August 2022.

5. Sho Aoki, Jared B Smith, Hao Li, Xunyi Yan, Masakazu Igarashi, Patrice Coulon, Jeffery R Wickens, Tom J H Ruigrok, and Xin Jin. An open cortico-basal ganglia loop allows limbic control over motor output via the nigrothalamic pathway. Elife, 8:e49995, September 2019.

6. Bernard W Balleine, Mauricio R Delgado, and Okihide Hikosaka. The role of the dorsal striatum in reward and decision-making. J. Neurosci., 27(31):8161–8165, August 2007.

7. Bilal A. Bari and Samuel J. Gershman. Undermatching is a consequence of policy compression. Journal of Neuroscience, 43(3):447–457, 2023. ISSN 0270-6474. doi: 10.1523/JNEUROSCI.1003-22.2022. URL https://www.jneurosci.org/content/43/3/447.

8. Jeff A. Beeler, Nathaniel Daw, Cristianne R. M. Frazier, and Xiaoxi Zhuang. Tonic dopamine modulates exploitation of reward learning. Frontiers in Behavioral Neuroscience, 4:170, 2010. doi: 10.3389/fnbeh.2010.00170.

9. Rafal Bogacz and Kevin Gurney. The basal ganglia and cortex implement optimal decision making between alternative actions. Neural Computation, 19(2):442–477, 2007. doi: 10.1162/neco.2007.19.2.442.

10. Anna Castañé, David E. H. Theobald, and Trevor W. Robbins. Selective lesions of the dorsomedial striatum impair serial spatial reversal learning in rats. Behavioural Brain Research, 210(1): 74–83, 2010. doi: 10.1016/j.bbr.2010.02.017.

11. Mark M. Churchland, John P. Cunningham, Matthew T. Kaufman, Justin D. Foster, Paul Nuyujukian, Stephen I. Ryu, and Krishna V. Shenoy. Neural population dynamics during reaching. Nature, 487(7405):51–56, 2012. doi: 10.1038/nature11129.

12. Nathaniel D Daw, Yael Niv, and Peter Dayan. Uncertainty-based competition between prefrontal and dorsolateral striatal systems for behavioral control. Nat. Neurosci., 8(12):1704–1711, December 2005.

13. Marco Del Giudice and Bernard J Crespi. Basic functional trade-offs in cognition: An integrative framework. Cognition, 179:56–70, October 2018.

14. Yue Du, John W. Krakauer, and Adrian M. Haith. The relationship between habits and motor skills in humans. Trends in Cognitive Sciences, 26(5):371–387, 2022.

15. Roy Fox, Ari Pakman, and Naftali Tishby. Taming the noise in reinforcement learning via soft updates. In Proceedings of the 32nd Conference on Uncertainty in Artificial Intelligence (UAI), 2016.

16. Michael J Frank. Computational models of motivated action selection in corticostriatal circuits. Current Opinion in Neurobiology, 21(3):381–386, 2011. ISSN 0959-4388. doi: 10.1016/j.conb.2011.02.013. URL https://www.sciencedirect.com/science/article/pii/S0959438811000407. Behavioural and cognitive neuroscience.

17. Alexandre Galashov, Siddhant M. Jayakumar, Leonard Hasenclever, Dhruva Tirumala, Jonathan Schwarz, Guillaume Desjardins, Wojciech M. Czarnecki, Yee Whye Teh, Razvan Pascanu, and Nicolas Heess. Information asymmetry in KL-regularized RL. In International Conference on Learning Representations (ICLR), 2019.

18. Matthieu Geist, Bruno Scherrer, and Olivier Pietquin. A theory of regularized Markov decision processes. In Proceedings of the 36th International Conference on Machine Learning, volume 97 of Proceedings of Machine Learning Research, pages 2160–2169. PMLR, 09–15 Jun 2019. URL https://proceedings.mlr.press/v97/geist19a.html.

19. Apostolos P. Georgopoulos, Andrew B. Schwartz, and Ronald E. Kettner. Neuronal population coding of movement direction. Science, 233(4771):1416–1419, 1986. doi: 10.1126/science. 3749885. URL https://www.science.org/doi/abs/10.1126/science.3749885.

20. Samuel J Gershman. Origin of perseveration in the trade-off between reward and complexity. Cognition, 204:104394, July 2020.

21. David H Gire, Vikrant Kapoor, Annie Arrighi-Allisan, Agnese Seminara, and Venkatesh N Murthy. Mice develop efficient strategies for foraging and navigation using complex natural stimuli. Curr. Biol., 26(10):1261–1273, May 2016.

22. Ann M Graybiel. Habits, rituals, and the evaluative brain. Annu. Rev. Neurosci., 31(1):359–387, July 2008.

23. Ann M Graybiel and Scott T Grafton. The striatum: where skills and habits meet. Cold Spring Harb. Perspect. Biol., 7(8):a021691, August 2015.

24. S N Haber, J L Fudge, and N R McFarland. Striatonigrostriatal pathways in primates form an ascending spiral from the shell to the dorsolateral striatum. J. Neurosci., 20(6):2369–2382, March 2000.

25. Adrian M Haith and John W Krakauer. The multiple effects of practice: skill, habit and reduced cognitive load. Curr Opin Behav Sci, 20:196–201, April 2018.

26. Arif A. Hamid, Jeffrey R. Pettibone, Omar S. Mabrouk, Vaughn L. Hetrick, Robert Schmidt, Caitlin M. Vander Weele, Robert T. Kennedy, Brandon J. Aragona, and Joshua D. Berke. Mesolimbic dopamine signals the value of work. Nature Neuroscience, 19(1):117–126, 2016. doi: 10.1038/nn.4173.

27. Alan N. Hampton, Peter Bossaerts, and John P. O’Doherty. The role of the ventromedial prefrontal cortex in abstract state-based inference during decision making in humans. Journal of Neuroscience, 26(32):8360–8367, 2006. doi: 10.1523/JNEUROSCI.1010-06.2006.

28. Houri Hintiryan, Nicholas N. Foster, Ian Bowman, Monica Bay, Muye Y. Song, Lin Gou, Seita Yamashita, Michael S. Bienkowski, Brian Zingg, Maxwell Zhu, X. William Yang, Jean C. Shih, Arthur W. Toga, and Hong-Wei Dong. The mouse cortico-striatal projectome. Nature Neuroscience, 19(8):1100–1114, 2016. doi: 10.1038/nn.4332.

29. Barbara J Hunnicutt, Bart C Jongbloets, William T Birdsong, Katrina J Gertz, Haining Zhong, and Tianyi Mao. A comprehensive excitatory input map of the striatum reveals novel functional organization. eLife, 5:e19103, nov 2016. ISSN 2050-084X. doi: 10.7554/eLife.19103. URL 10.7554/eLife.19103.

30. Daphna Joel, Yael Niv, and Eytan Ruppin. Actor–critic models of the basal ganglia: new anatomical and computational perspectives. Neural Networks, 15(4):535–547, 2002. ISSN 0893-6080. doi: 10.1016/S0893-6080(02)00047-3. URL https://www.sciencedirect.com/science/article/pii/S0893608002000473.

31. Risa Kawai, Timothy Markman, Rajesh Poddar, Raymond Ko, Antoniu L Fantana, Ashesh K Dhawale, Adam R Kampff, and Bence P Ölveczky. Motor cortex is required for learning but not for executing a motor skill. Neuron, 86(3):800–812, 2015.

32. Áron Kőszeghy, Wei Xu, Mingshan Liu, Peiheng Lu, Long Wan, Tong Deng, Sungmin Kang, Peggy Seriès, and Jian Gan. Medial prefrontal cortex activity precedes dorsomedial striatum in need for change during history-based flexible behavior. iScience, 28(12):113913, 2025. doi: 10.1016/j.isci.2025.113913.

33. David A Kupferschmidt, Konrad Juczewski, Guohong Cui, Kari A Johnson, and David M Lovinger. Parallel, but dissociable, processing in discrete corticostriatal inputs encodes skill learning. Neuron, 96(2):476–489.e5, October 2017.

34. Lucy Lai and Samuel J Gershman. Policy compression: An information bottleneck in action selection. In Psychology of Learning and Motivation. Academic Press, April 2021.

35. Lucy Lai and Samuel J Gershman. Human decision making balances reward maximization and policy compression. PLOS Computational Biology, 20(4):e1012057, 2024.

36. Lucy Lai, Ann Z. X. Huang, and Samuel J. Gershman. Action chunking as conditional policy compression. Cognition, 264:106201, 2025. doi: 10.1016/j.cognition.2025.106201.

37. Talia N. Lerner. Interfacing behavioral and neural circuit models for habit formation. Journal of Neuroscience Research, 98(6):1031–1045, 2020. doi: 10.1002/jnr.24581. URL https://onlinelibrary.wiley.com/doi/abs/10.1002/jnr.24581.

38. Sergey Levine. Reinforcement learning and control as probabilistic inference: Tutorial and review. *arXiv*, May 2018.

39. A. J. McGeorge and R. L. M. Faull. The organization of the projection from the cerebral cortex to the striatum in the rat. Neuroscience, 29(3):503–537, 1989.

40. John G Mikhael, Lucy Lai, and Samuel J Gershman. Rational inattention and tonic dopamine. *PLoS Comput. Biol.*, 17(3):e1008659, March 2021.

41. Earl K. Miller and Jonathan D. Cohen. An integrative theory of prefrontal cortex function. Annual Review of Neuroscience, 24:167–202, 2001.

42. Jonathan W Mink. The basal ganglia: focused selection and inhibition of competing motor programs. Prog. Neurobiol., 50(4):381–425, November 1996. doi: 10.1016/S0301-0082(96) 00042-1.

43. Kevin G C Mizes, Jack Lindsey, G Sean Escola, and Bence P Ölveczky. Dissociating the contributions of sensorimotor striatum to automatic and visually guided motor sequences. Nat. Neurosci., 26(10):1791–1804, October 2023.

44. Ali Mohebi, Jeffrey R. Pettibone, Arif A. Hamid, Jenny-Marie T. Wong, Leah T. Vinson, Tommaso Patriarchi, Lin Tian, Robert T. Kennedy, and Joshua D. Berke. Dissociable dopamine dynamics for learning and motivation. Nature, 570:65–70, 2019. doi: 10.1038/s41586-019-1235-y.

45. Ted Moskovitz, Kevin J. Miller, Maneesh Sahani, and Matthew M. Botvinick. Understanding dual process cognition via the minimum description length principle. PLOS Computational Biology, 20(10):e1012383, 2024.

46. Masayoshi Murakami, M Inês Vicente, Gil M Costa, and Zachary F Mainen. Neural antecedents of self-initiated actions in secondary motor cortex. Nature Neuroscience, 17(11):1574–1582, 2014.

47. Yael Niv, Daphna Joel, and Peter Dayan. A normative perspective on motivation. Trends in Cognitive Sciences, 10(8):375–381, 2006. doi: 10.1016/j.tics.2006.06.010.

48. Yael Niv, Nathaniel D Daw, Daphna Joel, and Peter Dayan. Tonic dopamine: opportunity costs and the control of response vigor. Psychopharmacology, 191(3):507–520, April 2007.

49. Heesun Park, Hoyoung Doh, Eunhwi Lee, Harhim Park, and Woo-Young Ahn. The neurocognitive role of working memory load when pavlovian motivational control affects instrumental learning. PLoS Comput. Biol., 19(12):e1011692, December 2023.

50. Naama Parush, Naftali Tishby, and Hagai Bergman. Dopaminergic balance between reward maximization and policy complexity. Front. Syst. Neurosci., 5:22, May 2011.

51. Payam Piray and Nathaniel D. Daw. Linear reinforcement learning in planning, grid fields, and cognitive control. Nature Communications, 12(1):4942, 2021.

52. Jonathan Rubin, Ohad Shamir, and Naftali Tishby. Trading value and information in MDPs. In Decision Making with Imperfect Decision Makers, pages 57–74. Springer Berlin Heidelberg, Berlin, Heidelberg, 2012.

53. John Schulman, Sergey Levine, Philipp Moritz, Michael I. Jordan, and Pieter Abbeel. Trust region policy optimization. In Proceedings of the 32nd International Conference on Machine Learning (ICML), volume 37 of PMLR, pages 1889–1897, 2015.

54. Susanne Still and Doina Precup. An information-theoretic approach to curiosity-driven reinforcement learning. Theory Biosci., 131(3):139–148, September 2012.

55. Richard S. Sutton and Andrew G. Barto. Reinforcement Learning: An Introduction. MIT Press, Cambridge, MA, 2nd edition, 2018.

56. Yee Whye Teh, Victor Bapst, Wojciech M. Czarnecki, John Quan, James Kirkpatrick, Raia Hadsell, Nicolas Heess, and Razvan Pascanu. Distral: Robust multitask reinforcement learning. In I. Guyon, U. Von Luxburg, S. Bengio, H. Wallach, R. Fergus, S. Vishwanathan, and R. Garnett, editors, Advances in Neural Information Processing Systems, volume 30. Curran Associates, Inc., 2017. URL https://proceedings.neurips.cc/paper_files/paper/2017/file/0abdc563a06105aee3c6136871c9f4d1-Paper.pdf.

57. Catherine A Thorn and Ann M Graybiel. Differential entrainment and learning-related dynamics of spike and local field potential activity in the sensorimotor and associative striatum. J. Neurosci., 34(8):2845–2859, February 2014.

58. Catherine A Thorn, Hisham Atallah, Mark Howe, and Ann M Graybiel. Differential dynamics of activity changes in dorsolateral and dorsomedial striatal loops during learning. Neuron, 66(5): 781–795, June 2010.

59. Dhruva Tirumala, Hyeonwoo Noh, Alexandre Galashov, Leonard Hasenclever, Arun Ahuja, Greg Wayne, Razvan Pascanu, Yee Whye Teh, and Nicolas Heess. Exploiting hierarchy for learning and transfer in KL-regularized RL. *arXiv*, March 2019.

60. Naftali Tishby and Daniel Polani. Information theory of decisions and actions. In Vassilis Cutsuridis, Amir Hussain, and John G. Taylor, editors, Perception-Action Cycle: Models, Architectures, and Hardware, pages 601–636. Springer, New York, NY, 2011.

61. Emanuel Todorov. Efficient computation of optimal actions. Proceedings of the National Academy of Sciences, 106(28):11478–11483, 2009. doi: 10.1073/pnas.0710743106. URL https://www.pnas.org/doi/abs/10.1073/pnas.0710743106.

62. Karly M Turner, Anna Svegborn, Mia Langguth, Colin McKenzie, and Trevor W Robbins. Opposing roles of the dorsolateral and dorsomedial striatum in the acquisition of skilled action sequencing in rats. J. Neurosci., 42(10):2039–2051, March 2022.

63. Henry H Yin and Barbara J Knowlton. The role of the basal ganglia in habit formation. Nat. Rev. Neurosci., 7(6):464–476, June 2006.

64. Henry H Yin, Barbara J Knowlton, and Bernard W Balleine. Lesions of dorsolateral striatum preserve outcome expectancy but disrupt habit formation in instrumental learning. Eur. J. Neurosci., 19(1):181–189, January 2004.

65. Henry H Yin, Sean B Ostlund, Barbara J Knowlton, and Bernard W Balleine. The role of the dorsomedial striatum in instrumental conditioning. Eur. J. Neurosci., 22(2):513–523, July 2005.

66. Hong Yu, Xinkuan Xiang, Zongming Chen, Xu Wang, Jiaqi Dai, Xinxin Wang, Pengcheng Huang, Zheng-Dong Zhao, Wei L Shen, and Haohong Li. Periaqueductal gray neurons encode the sequential motor program in hunting behavior of mice. Nat. Commun., 12(1):6523, November 2021.

