## Supplementary Figures S1-S8 for "Policy regularization as a unifying theory of the striatal division of labor in learning"

### Parameter robustness sweeps

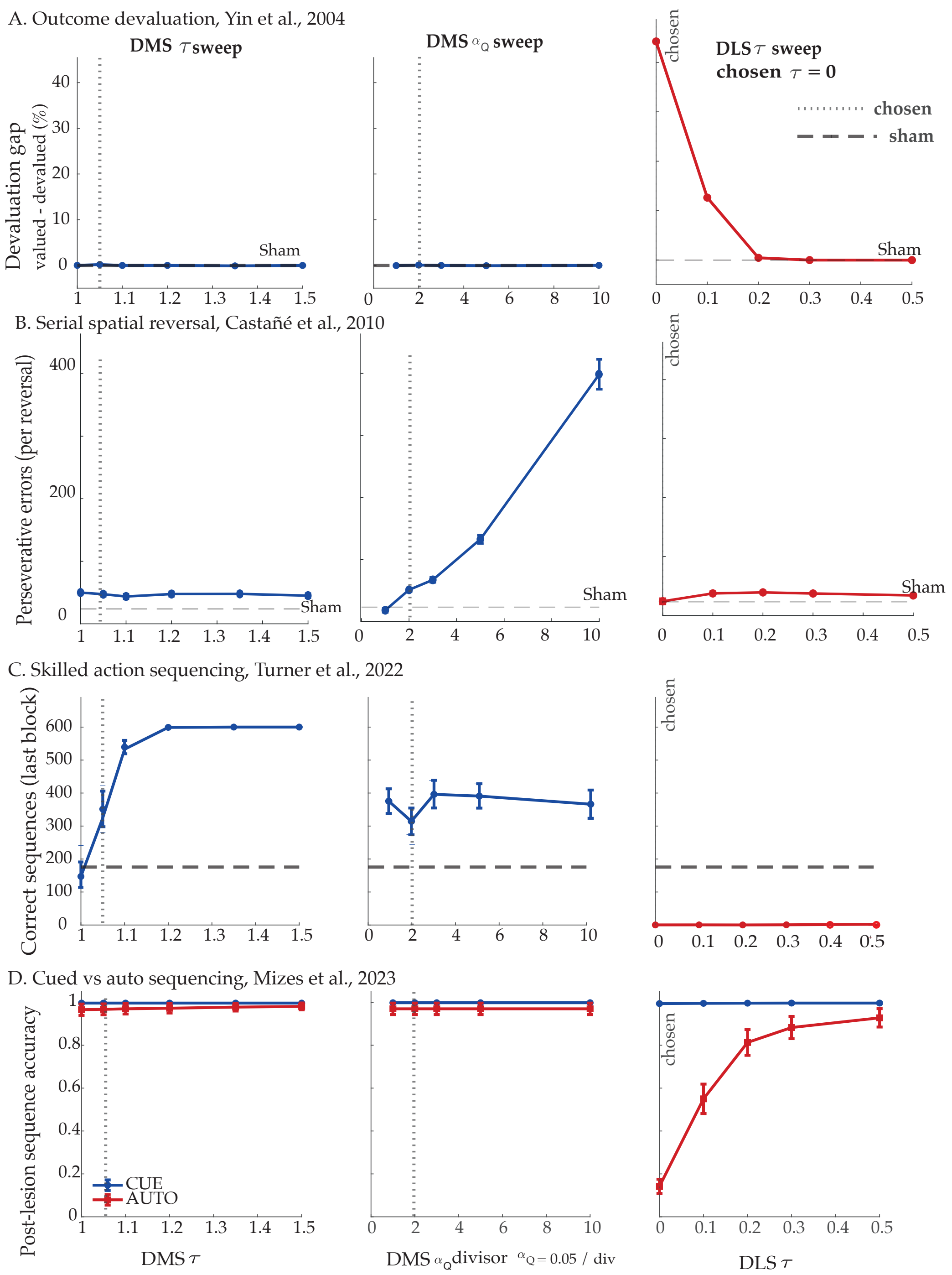

**Figure S1. Robustness of dissociations to the lesion parameters.** Each row is one paradigm; each column sweeps a single lesion parameter (DMS $\tau$, DMS $\alpha_Q$ divisor, DLS $\tau$) while holding the other parameters constant at their default value. The dotted line is the chosen value used in the article (DMS lesion $\tau = 1.05$ and $\alpha_Q = \alpha_Q/2$; DLS lesion $\tau = 0$, the leftmost point); dashed line: Sham baseline. Line color distinguishes the lesion (DMS, blue; DLS, red); $n = 25$ agents per point, mean ± SEM. In (D), blue and red denote the CUE and AUTO tasks. **(A) Outcome devaluation (Yin et al., 2004):** the devaluation gap (valued − devalued) stays at Sham level ($\approx 0$) across both DMS parameters. DLS lesions produce devaluation sensitivity only when near-complete: the gap is largest at $\tau = 0$ and returns to Sham level by $\tau \approx 0.2$. **(B) Serial spatial reversal (Castañé et al., 2010):** perseverative errors per reversal increase monotonically with the $\alpha_Q$ divisor and are flat across DMS $\tau$. Perseveration stays near Sham level across DLS $\tau$. **(C) Skilled action sequencing (Turner et al., 2022):** correct sequences in the last block rise with DMS $\tau$, from Sham level at $\tau = 1$ to ceiling by $\tau \approx 1.2$, and are flat across the $\alpha_Q$ divisor. DLS-lesioned agents stay at floor across $\tau = 0$–$0.5$ (see Figure S8 for the full range). **(D) Cued vs. automatic sequencing (Mizes et al., 2023):** DMS lesions spare both cued and automatic execution at all values. DLS lesions spare cued execution; automatic execution is impaired at $\tau = 0$ and recovers as $\tau$ rises, approaching pre-lesion accuracy by $\tau = 0.5$, a value at which the five-step sequence in (C) remains at floor. Because slowing $\alpha_Q$ is required for the reversal deficit (B) and raising $\tau$ for enhanced sequencing (C), each DMS manipulation is necessary for a different paradigm.

### Policy regularization vs Vanilla RL

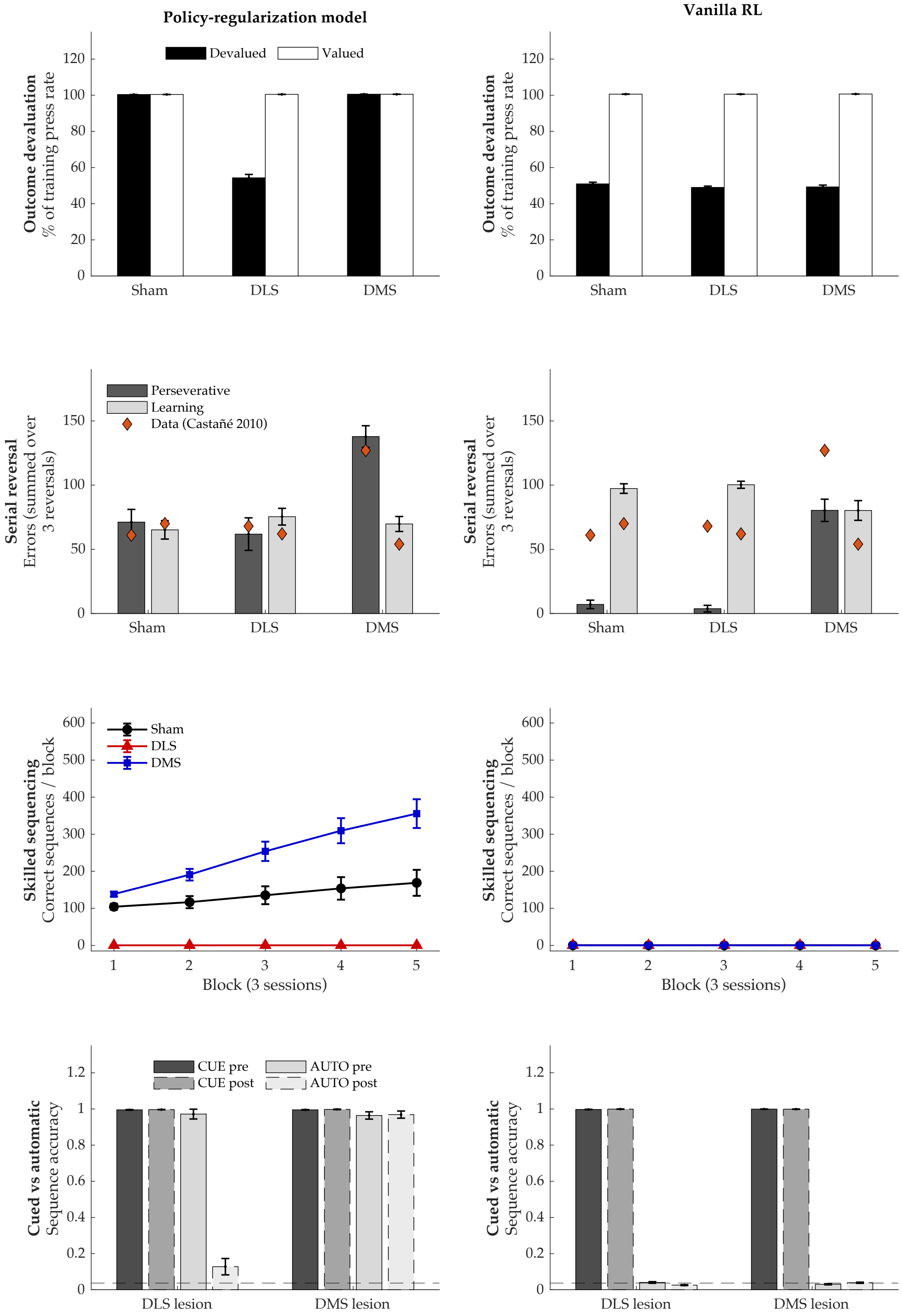

**Figure S2. Comparison of the policy-regularization model with vanilla reinforcement learning (no default policy).** Each row pairs our policy-regularization model (left) with a vanilla soft-Q-learning control (right; $\tau = 0$ throughout, removing $\pi_0$ from both the policy and the value update); $y$-axes are shared within each row. $n = 25$ agents per group. **(A) Outcome devaluation:** our model dissociates (DLS sensitive; DMS and Sham habitual), whereas vanilla RL shows devaluation sensitivity in all groups (no habit). **(B) Serial spatial reversal (errors summed across three reversals):** both models produce elevated DMS perseveration, because this effect depends on slow updating of $Q$, which vanilla RL retains. Without $\pi_0$, however, Sham and DLS-lesioned agents almost never perseverate (2.4 and 1.3 errors per reversal, vs. 23.7 and 20.6 in the full model), unlike animals in all groups. **(C) Skilled action sequencing:** our model reproduces the DMS > Sham > DLS ordering, whereas vanilla RL performs at $\approx 0$ in all groups, because there is no cached chain to drive uncued execution. **(D) Cued vs. automatic sequencing:** vanilla RL executes cued sequences, but automatic execution is at chance (dashed line) before and after the lesion in both groups, so automaticity never forms. Thus $\pi_0$ is required for the habit (A) and automaticity (C, D) dissociations, and for the baseline perseveration of intact animals (B).

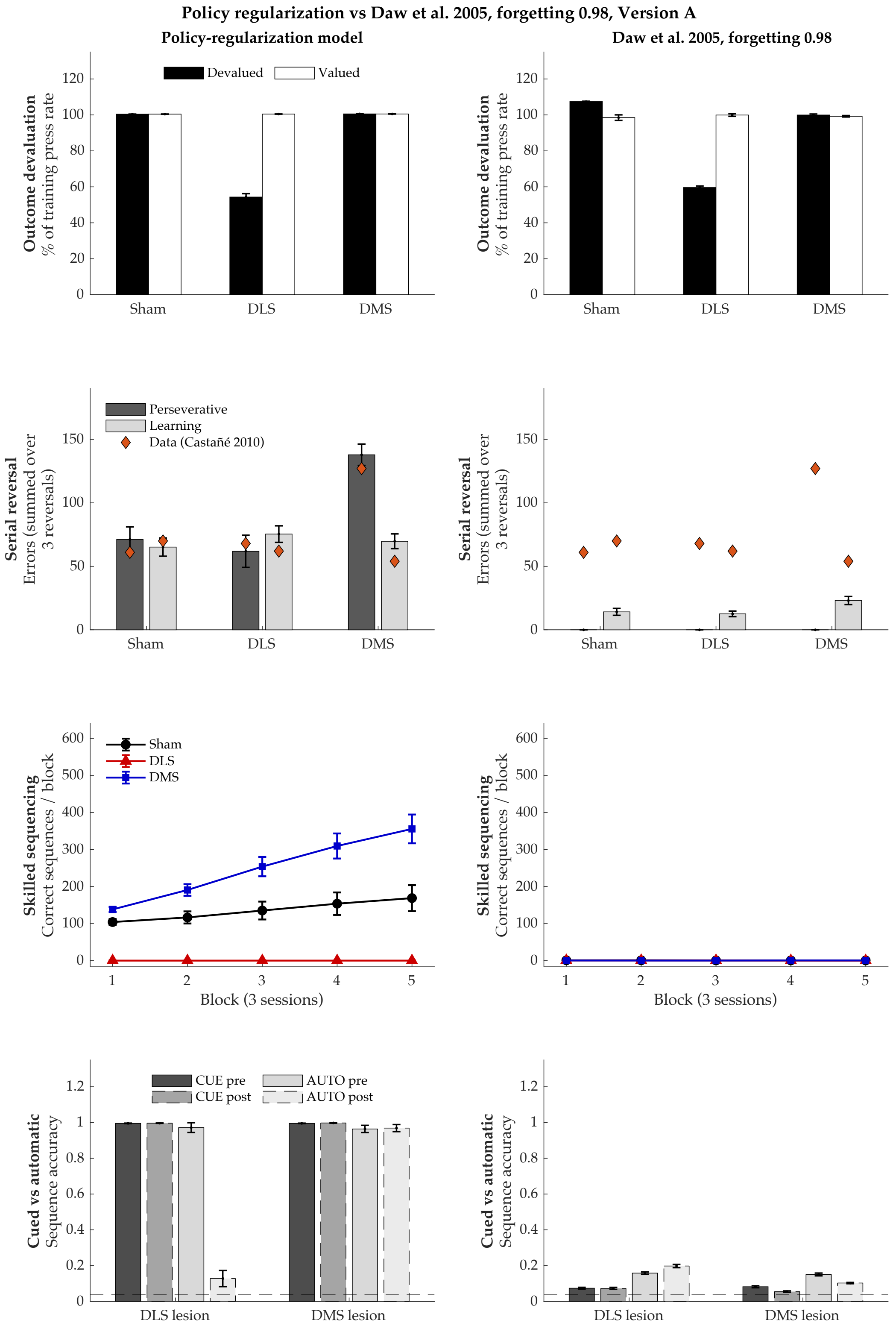
Policy regularization vs Daw et al. 2005, forgetting 0.98, Version A
Policy-regularization model
Daw et al. 2005, forgetting 0.98
Devalued
Valued
Outcome devaluation
% of training press rate
Sham
DLS
DMS
Perseverative
Learning
Data (Castañé 2010)
Serial reversal
Errors (summed over 3 reversals)
Sham
DLS
DMS
Skilled sequencing
Correct sequences / block
Block (3 sessions)
CUE pre
CUE post
AUTO pre
AUTO post
Cued vs automatic
Sequence accuracy
DLS lesion
DMS lesion

**Figure S3. Policy regularization model vs. the dual-controller model of Daw et al. (2005): forgetting factor 0.98, sequence position known.** Left: policy-regularization model. Right: dual-controller model, with a model-free (cached) and a model-based (tree-search) controller, simulated with the published parameters, including the published forgetting factor (0.98), on the same paradigms, training protocols and lesion timings. Both controllers learn throughout; a DLS lesion removes the cached controller and a DMS lesion removes the tree-search controller from action selection. In the uncued sequence tasks (C, D), the state given to both controllers includes the agent's current position in the sequence. $n$ = 25 agents per group, mean ± SEM; diamonds in (B): data from Castañé et al. (2010); dashed line in (D): chance. **(A) Outcome devaluation:** the dual-controller model reproduces the dissociation. After a DLS lesion only tree search remains, and it uses the devalued outcome immediately, so pressing decreases; the cached controller updates the value of pressing only after experiencing the devalued outcome, so Sham and DMS-lesioned agents continue to press. **(B) Serial spatial reversal:** no group makes perseverative errors. Forgetting returns all learned quantities toward their priors at every timestep, so the previously rewarded lever is forgotten before each reversal. **(C) Skilled action sequencing:** no group learns the sequence. With forgetting at every timestep, the model retains only the most recent ≈ 50 timesteps of experience, too few for the reward delivered after the fifth poke to build up value for the earlier pokes. **(D) Cued vs. automatic sequencing:** for the same reason, neither cued nor automatic sequences are learned, before or after either lesion.

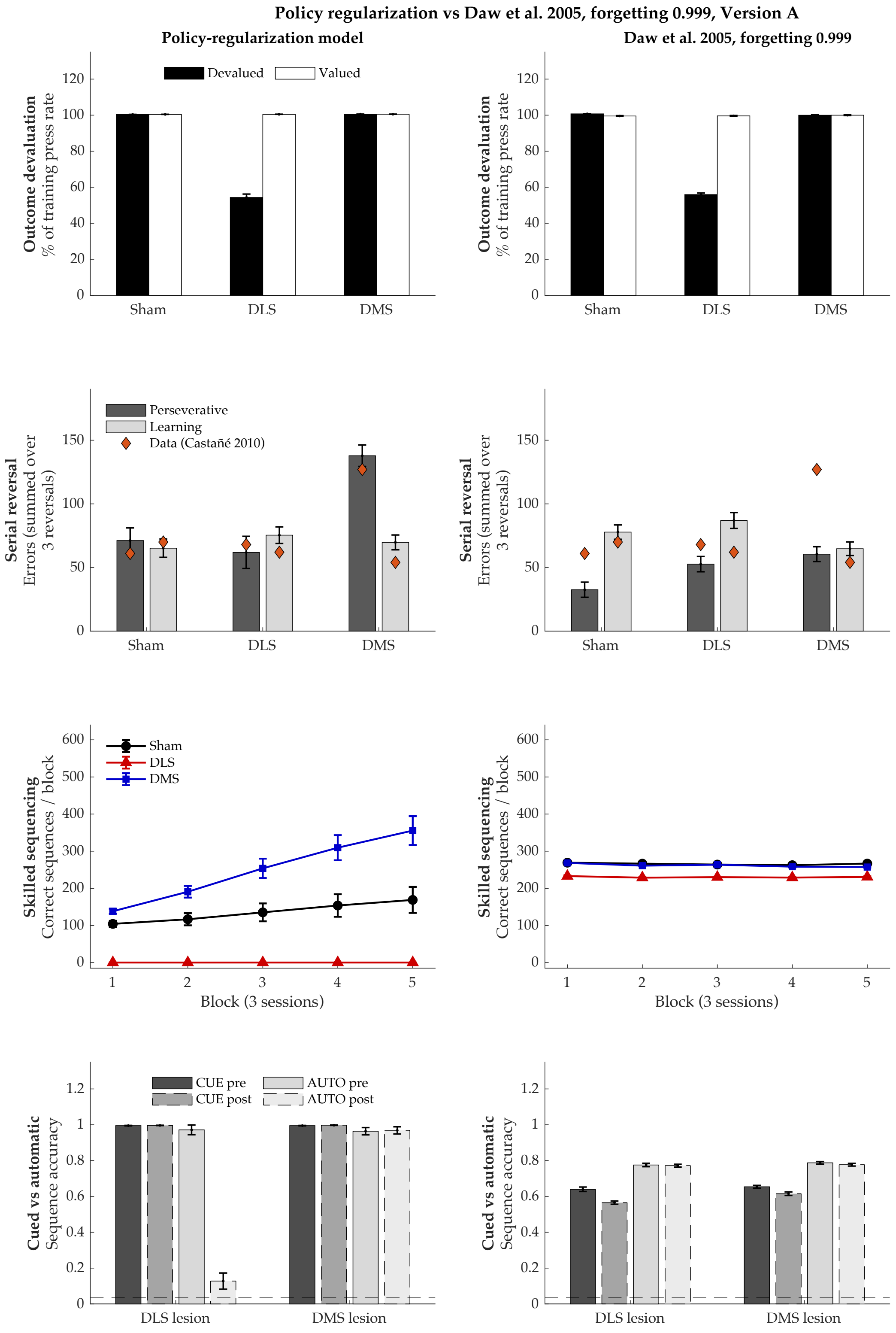

Policy regularization vs Daw et al. 2005, forgetting 0.999, Version A
Policy-regularization model
Daw et al. 2005, forgetting 0.999
Outcome devaluation
% of training press rate
Devalued
Valued
Sham
DLS
DMS
Serial reversal
Errors (summed over 3 reversals)
Perseverative
Learning
Data (Castañé 2010)
Skilled sequencing
Correct sequences / block
Block (3 sessions)
Cued vs automatic
Sequence accuracy
CUE pre
CUE post
AUTO pre
AUTO post
DLS lesion
DMS lesion

**Figure S4. Comparison with the dual-controller model of Daw et al. (2005): forgetting factor 0.999, sequence position known.** As in Figure S3, with slower forgetting (0.999), so that learning persists across sessions. **(A) Outcome devaluation:** the dissociation is reproduced, for the same reason as in Figure S3. **(B) Serial spatial reversal:** both DMS- and DLS-lesioned agents make more perseverative errors than Sham, whereas in the data only DMS lesions do. Because both controllers learn the lever–reward contingency from the same experience, each retains the previous contingency after a reversal, and removing either one increases perseveration. **(C) Skilled action sequencing:** the sequence is learned, but neither lesion reproduces the data: DLS-lesioned agents perform only slightly below Sham, and DMS-lesioned agents do not exceed Sham. Because both controllers learn values over the same position states, either one can execute the sequence alone, so removing one has little effect. **(D) Cued vs. automatic sequencing:** for the same reason, DLS lesions do not impair automatic execution: tree search alone can execute the sequence from the position state.

**Policy regularization vs. Daw et al. (2005), version B, forgetting 0.98**

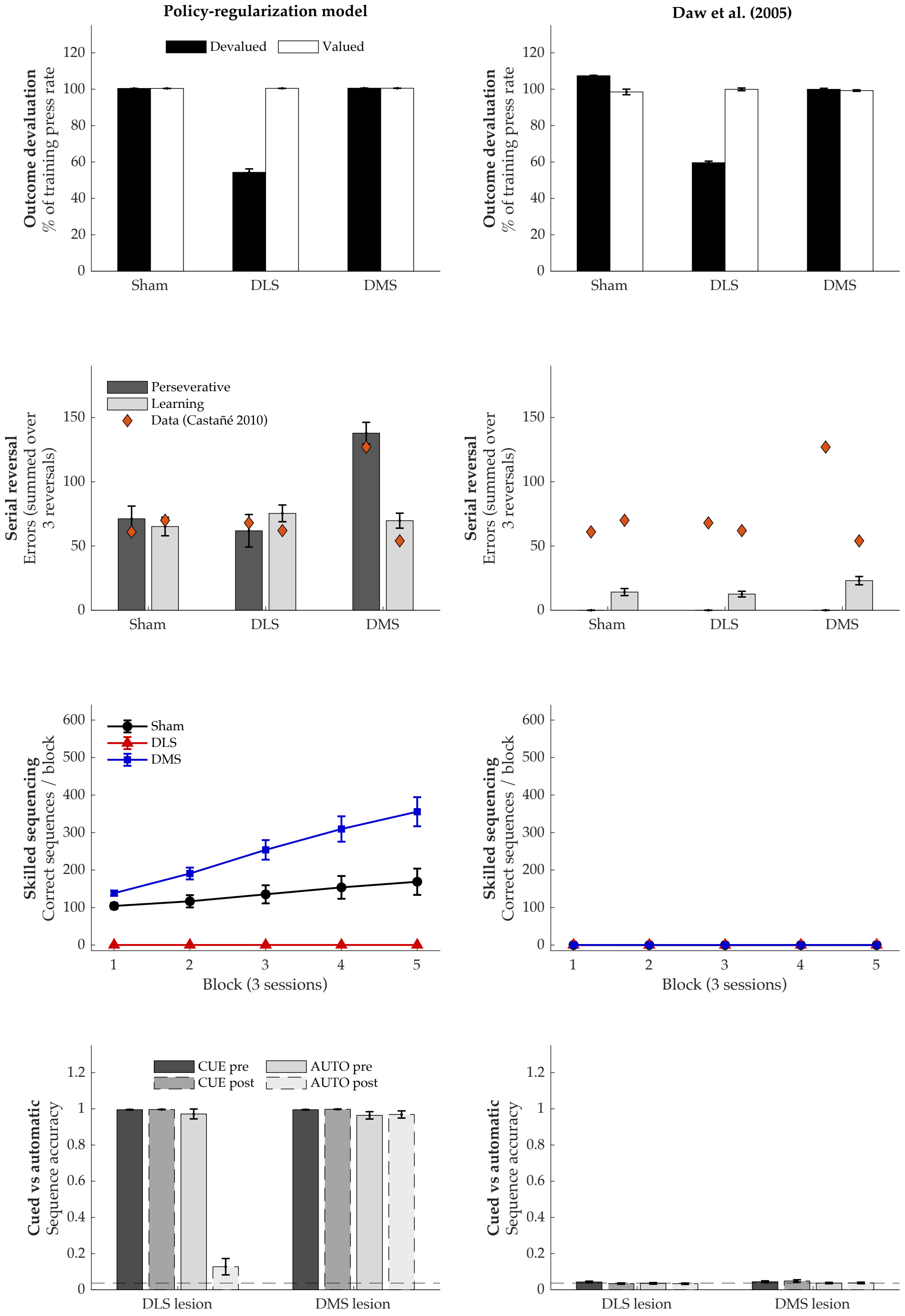

**Figure S5. Comparison with the dual-controller model of Daw et al. (2005): forgetting factor 0.98, sequence position unknown.** As in Figure S3, except that in the uncued sequence tasks the agent is not given its position in the sequence and acts from a uniform belief over the cue states. Panels (A) and (B) contain no uncued sequences and are identical to Figure S3. **(C, D) Sequencing:** no group performs above chance, before or after either lesion. Without cues, every press occurs in the same belief state, so neither controller can determine which action comes next.

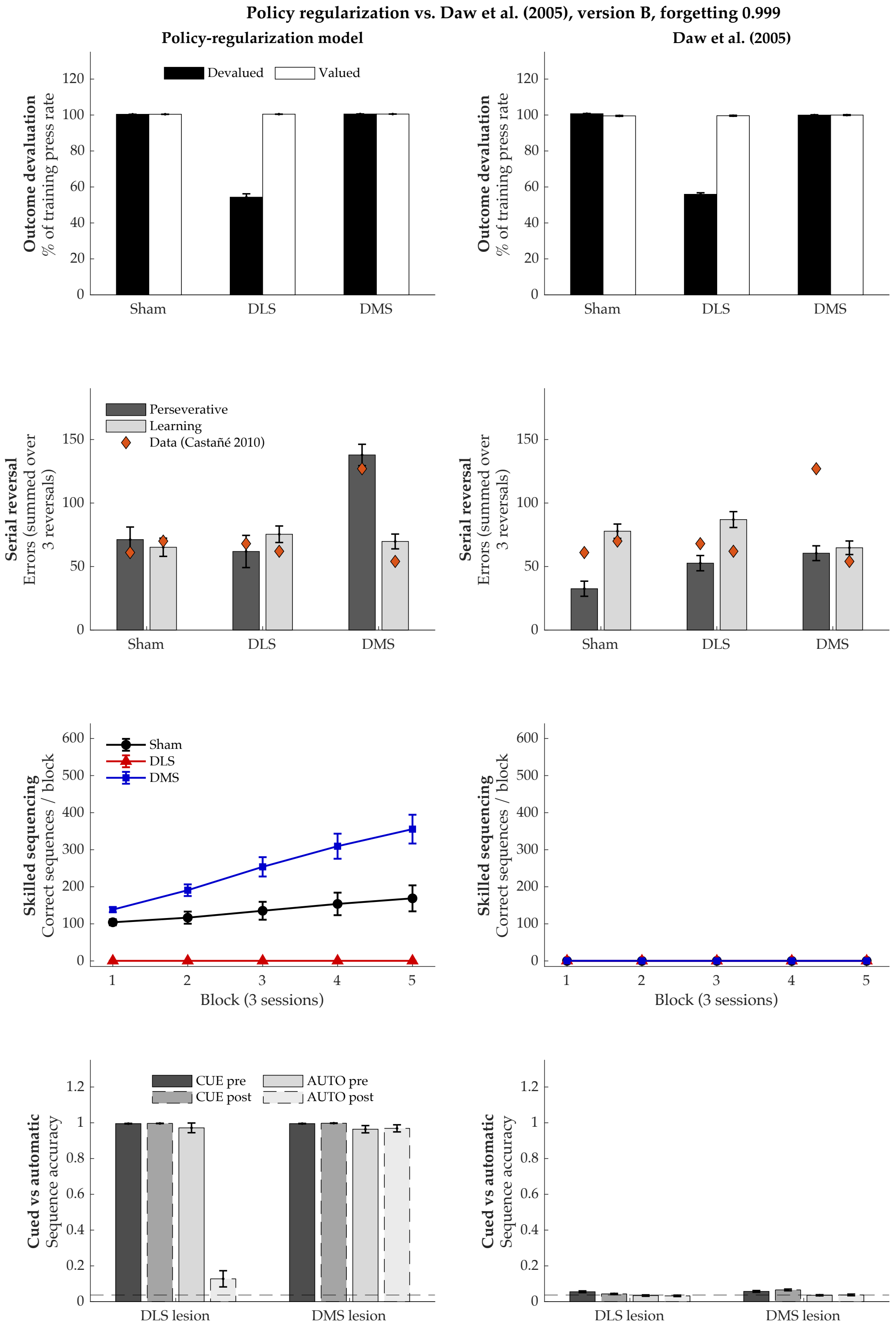
Policy regularization vs. Daw et al. (2005), version B, forgetting 0.999
Policy-regularization model
Daw et al. (2005)
Devalued
Valued
Outcome devaluation
% of training press rate
Sham
DLS
DMS
Perseverative
Learning
Data (Castañé 2010)
Serial reversal
Errors (summed over 3 reversals)
Skilled sequencing
Correct sequences / block
Block (3 sessions)
CUE pre
CUE post
AUTO pre
AUTO post
Cued vs automatic
Sequence accuracy
DLS lesion
DMS lesion

**Figure S6. Comparison with the dual-controller model of Daw et al. (2005): forgetting factor 0.999, sequence position unknown.** As in Figure S5, with forgetting factor 0.999. Panels (A) and (B) are identical to Figure S4. **(C, D) Sequencing:** even with slow forgetting, no group performs above chance. Both controllers select actions based on the current state; without cues or position information, every press occurs in the same state, so neither controller carries the sequential structure. In our model, by contrast, the default policy conditions on the previous action and can execute the sequence without cues.

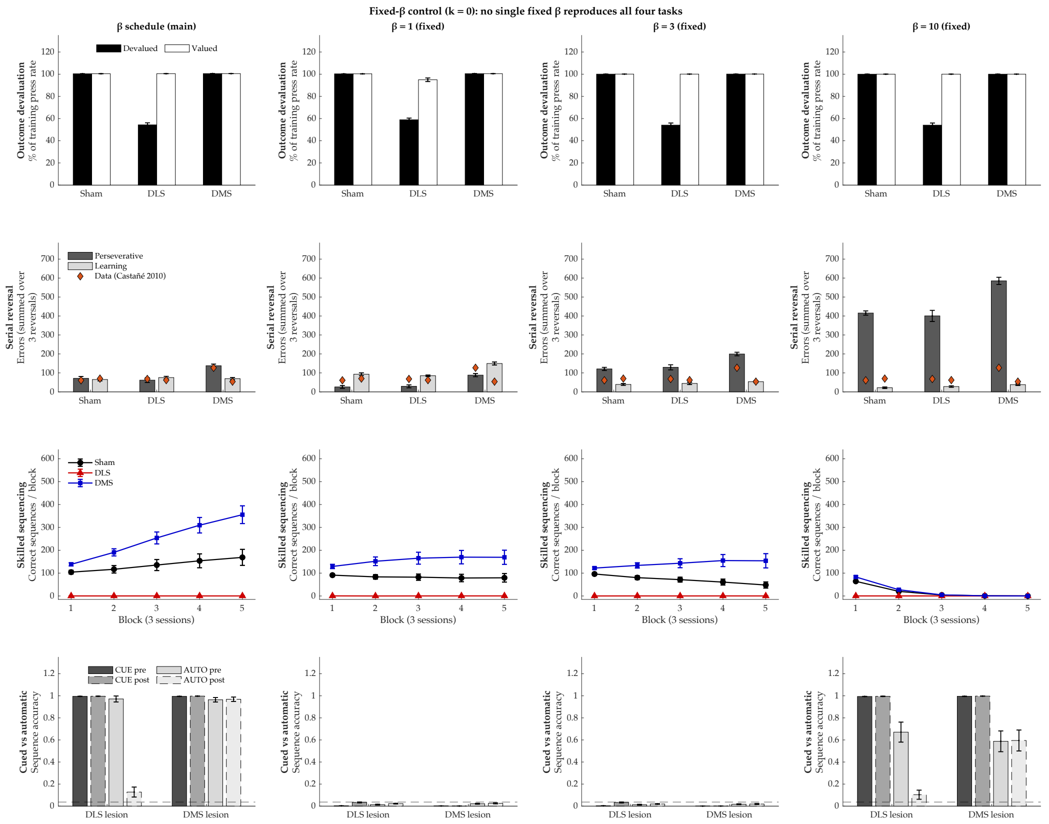


**Figure S7. No single fixed $\beta$ reproduces all four paradigms.** Columns: the $\beta$ schedule used in the main simulations (left), and $\beta$ held fixed at 1, 3 and 10 throughout ($k = 0$). All other parameters, training protocols and lesion timings are unchanged. $n = 25$ agents per group, mean ± SEM; diamonds in (B): data from Castañé et al. (2010); dashed line in (D): chance. **(A) Outcome devaluation:** the dissociation is reproduced at every $\beta$. **(B) Serial spatial reversal (errors summed across three reversals):** DMS-specific perseveration is present at every $\beta$, but its magnitude grows with $\beta$; at $\beta = 10$, all groups make roughly six times more perseverative errors than with the schedule, far above the data. **(C) Skilled action sequencing:** at $\beta = 1$ and 3, the DMS > Sham > DLS ordering is preserved, although Sham performance declines across blocks; at $\beta = 10$, all groups fall to $\approx 0$. **(D) Cued vs. automatic sequencing:** at $\beta = 1$ and 3, neither cued nor automatic sequences are learned (accuracy near chance); at $\beta = 10$, cued sequences are learned, but automatic execution is unreliable even before the lesion.

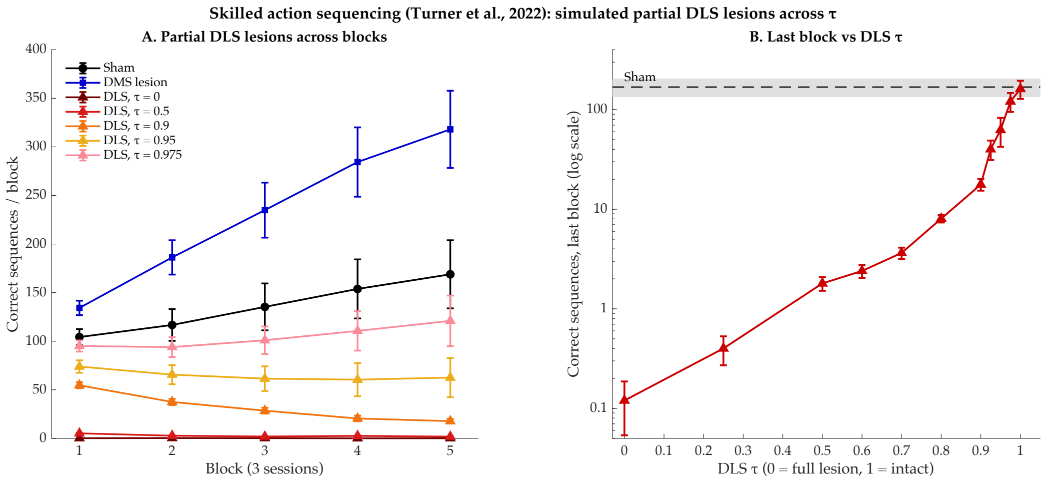


**Figure S8. Partial DLS lesions in skilled action sequencing (Turner et al., 2022).** **(A)** Correct sequences per block after the lesion for Sham, DMS-lesioned agents, and DLS lesions of decreasing severity ($\tau$ = 0, 0.5, 0.9, 0.95, 0.975; $\tau$ = 1 is intact). **(B)** Correct sequences in the last block as a function of DLS $\tau$ (log scale); dashed line and shading: Sham mean ± SEM. Because $\pi_0$ supplies each of the five transitions, a reduction in $\tau$ compounds across steps, giving a steep, nonlinear dependence on lesion size. $n$ = 25 agents per group, mean ± SEM.

### References


Anna Castañé, David E. H. Theobald, and Trevor W. Robbins. Selective lesions of the dorsomedial striatum impair serial spatial reversal learning in rats. *Behavioural Brain Research*, 210(1):74–83, 2010. doi: 10.1016/j.bbr.2010.02.017.

Nathaniel D Daw, Yael Niv, and Peter Dayan. Uncertainty-based competition between prefrontal and dorsolateral striatal systems for behavioral control. *Nat. Neurosci.*, 8(12):1704–1711, December 2005.

Kevin G C Mizes, Jack Lindsey, G Sean Escola, and Bence P Ölveczky. Dissociating the contributions of sensorimotor striatum to automatic and visually guided motor sequences. *Nat. Neurosci.*, 26(10):1791–1804, October 2023.

Karly M Turner, Anna Svegborn, Mia Langguth, Colin McKenzie, and Trevor W Robbins. Opposing roles of the dorsolateral and dorsomedial striatum in the acquisition of skilled action sequencing in rats. *J. Neurosci.*, 42(10):2039–2051, March 2022.

Henry H Yin, Barbara J Knowlton, and Bernard W Balleine. Lesions of dorsolateral striatum preserve outcome expectancy but disrupt habit formation in instrumental learning. *Eur. J. Neurosci.*, 19(1):181–189, January 2004.
